# A study of different pre-treatment methods for cigarettes and their aroma differences

**DOI:** 10.1101/2025.03.17.643679

**Authors:** Dengke Li, Yuchen Gu, Chuntao Zhang, Ting Fei, Lichao Ma, Ruoxin Wu, Liqi Tao, Zhizhang Tian, Tao Feng

## Abstract

The study innovatively utilized a combination of smoking machine and Cambridge filter to effectively extract the aroma compounds generated during smoking of cigarettes, and employed three different pretreatment methods to obtain optimal cigarette pretreatment procedures. In addition, the aroma components of different cigarettes were identified by different instruments such as gas chromatography-mass spectrometry. Combined with the calculation of Odor Activity Values (OAV), PCA analysis and sensory evaluation were performed on the five types of cigarettes to reveal differences and similarities in their aroma profiles.

## Introduction

Tobacco smoking has been a prevalent habit across cultures and societies for centuries, manifesting itself in various forms, with the tobacco cigarette being the most ubiquitous [1]. The extensive history and widespread usage of traditional tobacco cigarettes have prompted considerable scientific inquiry into their health implications, primarily focusing on the detrimental effects they pose to human health [2-9]. However, amidst this research landscape, there exists a noticeable disparity in the attention devoted to the analysis of aromatic compounds present in tobacco cigarettes. Conversely, investigations into the aromatic profile of electronic cigarettes have garnered substantial interest, surpassing that of traditional tobacco cigarettes [10-13]. The aromatic constituents of tobacco products play a pivotal role in shaping users’ sensory experiences and may contribute significantly to the overall appeal and addictive potential of these products. Despite their importance, the scientific exploration of these aromatic compounds, particularly within the context of traditional tobacco cigarettes, remains relatively limited [14]. This gap in understanding poses a critical barrier to comprehensively assessing the sensory and potentially addictive properties of tobacco cigarettes.

In light of this, the present study seeks to address this gap by undertaking a systematic analysis of the aromatic composition of five different variants of the same brand of tobacco cigarettes. By elucidating the specific aromatic compounds present in these cigarettes, we aim to enhance our understanding of the sensory attributes associated with traditional tobacco smoking. Furthermore, we endeavor to identify the most effective preprocessing methods for extracting and analyzing these aromatic compounds, thereby refining existing methodologies in tobacco research. To ensure the accuracy and reliability of our findings, we employ a novel experimental approach that eschews traditional chemical analysis of tobacco constituents. Instead, we utilize a smoking machine to simulate the act of cigarette smoking, replicating the inhalation process typically undertaken by human users [15, 16]. This methodological innovation allows for the precise capture and extraction of aromatic compounds formed during smoking, thereby providing a more nuanced insight into the composition of tobacco cigarette emissions.

Through the endeavors of this study, we aspire to shed light on the intricate relationship between the aromatic profile of tobacco cigarettes and their sensory characteristics. By advancing our understanding in this domain, we hope to inform public health initiatives and regulatory policies aimed at mitigating the harms associated with tobacco smoking, while simultaneously providing valuable insights into the development of safer and less addictive alternatives.

## Materials and Methods

### Samples and Instruments

Five different cigarettes from the same brand are yk, ku, jy, hd and lj; solid-phase extraction column Cleanert PEP-SPE (1000 mg packing): Agela & Phenomenex, Inc. Solvent-assisted flavor evaporation (SAFE): Glasblaserei-Bahr; Agilent 6890A gas chromatograph (GC) system with a 5975C mass spectrometer (MS): Agilent Technologies, Inc.; DVB/CAR/PDMS headspace solid-phase microextraction head: Supelco®; HP-INNOWax column (60 m/0.25 mm/0.25 μm): J&W Scientifc, Inc.

### Cambridge Filter Method

The PUFFMAN X200AF is a smoking machine designed to simulate the process of human smoking and consists of a variety of trapping systems that can be used for routine analyses, product development, and special analyses. The core structure of the PUFFMAN X200AF is the smoke trapping system, and in this experiment, a Cambridge filter was used to trap the smoke on the filter.

Before the experiment, the cigarettes and Cambridge filters were placed in a constant temperature of 26 LJ and a constant humidity of 60 % for 2 days, and the smoking parameters of the smoking machine were adopted from the ISO 3308 standard, with a suction volume of 35 mL/time, each time for 2 s, to collect the particle phase of the mainstream cigarette smoke, and to calculate the total particulate matter (TPM).

### Solid-Phase Microextraction (SPME)

Solid-phase microextraction (SPME) is a modern, nonexhaustive sample preparation technique. The advantages of this technique include simplicity, rapidity, improved sample clean-up, accurate analysis, and low organic solvent consumption [17]. A piece of Cambridge filter with total particulate phase material (TPM) mass of about 0.2 g was cut into pieces and packed into a headspace bottle, the extraction bottle with the sample was equilibrated in a water bath at 60 °C for 40 min. The sample was then extracted with a SPME extraction head (DVB/CAR/PDMS fiber) inserted into the extraction bottle at the same temperature for 40 min. After extraction, the fibers were transferred to the injection port and desorbed at 250 °C for 3 min for GC-MS analysis.

### Solvent-Assisted Flavor Evaporation (SAFE)

Solvent-assisted favor evaporation (SAFE) allows the isolation of volatiles from either solvent extracts, aqueous foods, aqueous food suspensions, or even matrices with a high oil content [18]. A piece of Cambridge filter tablet with the mass of total particulate phase material (TPM) of about 0.2 g was cut up and mounted in a 500 mL conical flask, 150 mL of methylene chloride was added, and the extraction was carried out for 1 h on a magnetic stirrer, then filtered and transferred to a separatory funnel, and the upper layer of the liquid was put into a 500 mL round-bottomed flask, and the lower layer of the liquid was retained and transferred to a conical flask, and the extraction was carried out for the second time with the addition of 150 mL methylene chloride, and the upper layer of liquid was transferred to a round-bottomed flask again after the third transfer to a conical flask. Extraction was carried out for 1 h on a magnetic stirrer, filtered again, transferred to a dispensing funnel, the upper layer of liquid was transferred to a round-bottomed flask again, the lower layer of liquid was retained, and the third time it was transferred to a conical flask, 100 mL of dichloromethane was added, and the upper layers of the three times were combined in a round-bottomed flask. Three sequential 1-hour extractions under magnetic stirring (total 3 hours). The fractions were dried with anhydrous sodium sulfate (5 g) and concentrated on a rotary evaporator to 5 mL, then further concentrated under a gentle stream of nitrogen to 1 mL. The concentrated fractions were stored at -20 °C prior to GC-MS analysis.

### Simultaneous Distillation Extraction (SDE)

When SDE is used correctly, it is often the best method for obtaining the highest recovery from a wide range of compounds [19]. A Cambridge filter piece of total particulate phase material (TPM) mass of about 0.2 g was cut up and added to a 500 mL flask with 100 mL of deionized water, and 50 mL of methylene chloride was added to a 150 mL flask. The SDE apparatus was assembled, and the 500 mL and 50 mL flasks were heated at the same time for 2 h at 100±1°C and 50±1°C, respectively, and then using the same method as the SAFE treatment, the extracted dichloromethane solution was dried, concentrated and stored.

### GC-MS Conditions

The GC column was equipped with HP-INNOWax (60 m×0.25 mm, 0.25 µm), and the carrier gas was helium with a flow rate of 1 mL/min. The injection volume was 1 μL with the split ratio 10:1. The programmed ramp-up procedure: the starting temperature was 40 LJ, held for 3 min; the temperature was ramped up to 150 LJ at 10 LJ/min, and then increased to 230 LJ at 3 LJ/min, and held for 5 min. The temperature of both the inlet and the auxiliary heating zone was 250°C. The samples were separated by the column and then entered the mass spectrometry for detection. The ion source was an electron ionization (EI) source with an electron energy of 70 eV and a mass scan range of 30–350 m/z. The temperature of the mass spectrometry interface was 250°C and the ion source temperature was 230°C; interface temperature 150 LJ; solvent delay time 5 min; multiplier voltage 1964.7 V, scanning range 20∼400 amu.

### Odor Activity Values (OAV)

The OAV represents the olfactory activity value of the favor active compound, OAV >1 means that the compound has a direct effect on the aroma, and it is generally accepted that the higher the OAV the greater the contribution of the volatile compound to the aroma of the sample. The calculation formula is as follows:

OAV=c/T

where c is the concentration of the compound, mg/kg; T is the threshold of the compound in water, mg/kg.

### Statistical Analysis

The NIST20 database was used for GC-MS analysis. The concentration of volatiles was submitted to analysis of variance (ANOVA) by SPSS version 26.0. Microsoft Office Excel 2021 was used for data processing and radar plotting. Principal component analysis (PCA) was performed using the Dynamic PCA plug-in to evaluate the regularity and difference among tested samples.

### Sensory Evaluation

In order to objectively, accurately and scientifically describe the smoke comfort of cigarette products, various evaluation methods in cigarette, food, beverage, wine, fragrance and other related industries at home and abroad were first studied and used for reference. Based on the theory of flavor and aroma, the research and practice of cigarette smoke comfort characteristics were carried out. Based on “YC/T 497-2014 Chinese cigarette style Sensory Evaluation Method” and “YC/T 496-2014 cigarette sensory Comfort Evaluation Method”, the sensory evaluation of cigarette smoking comfort was developed.

**Table 1.**
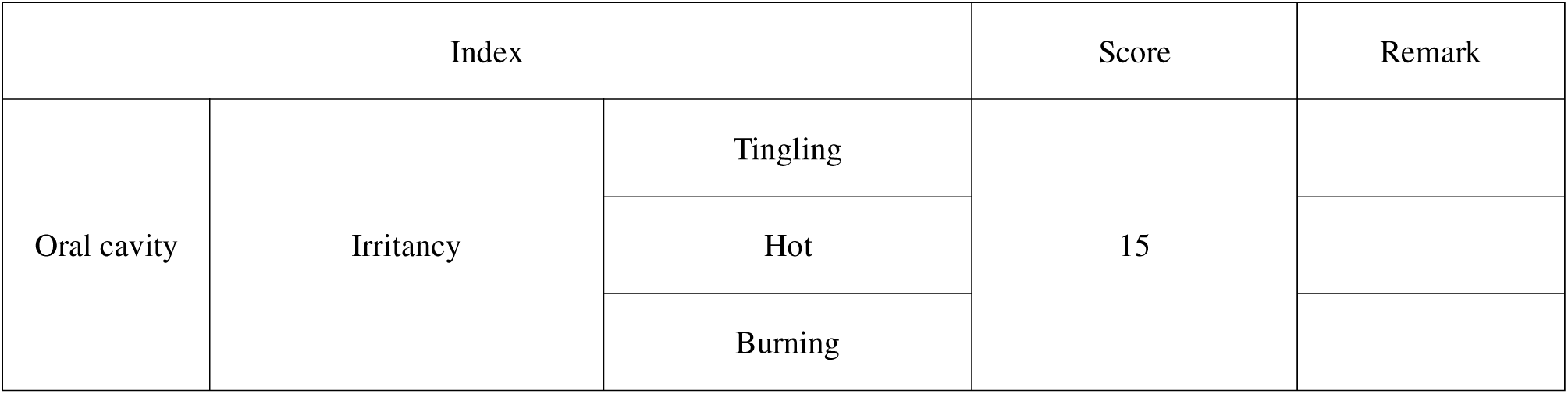
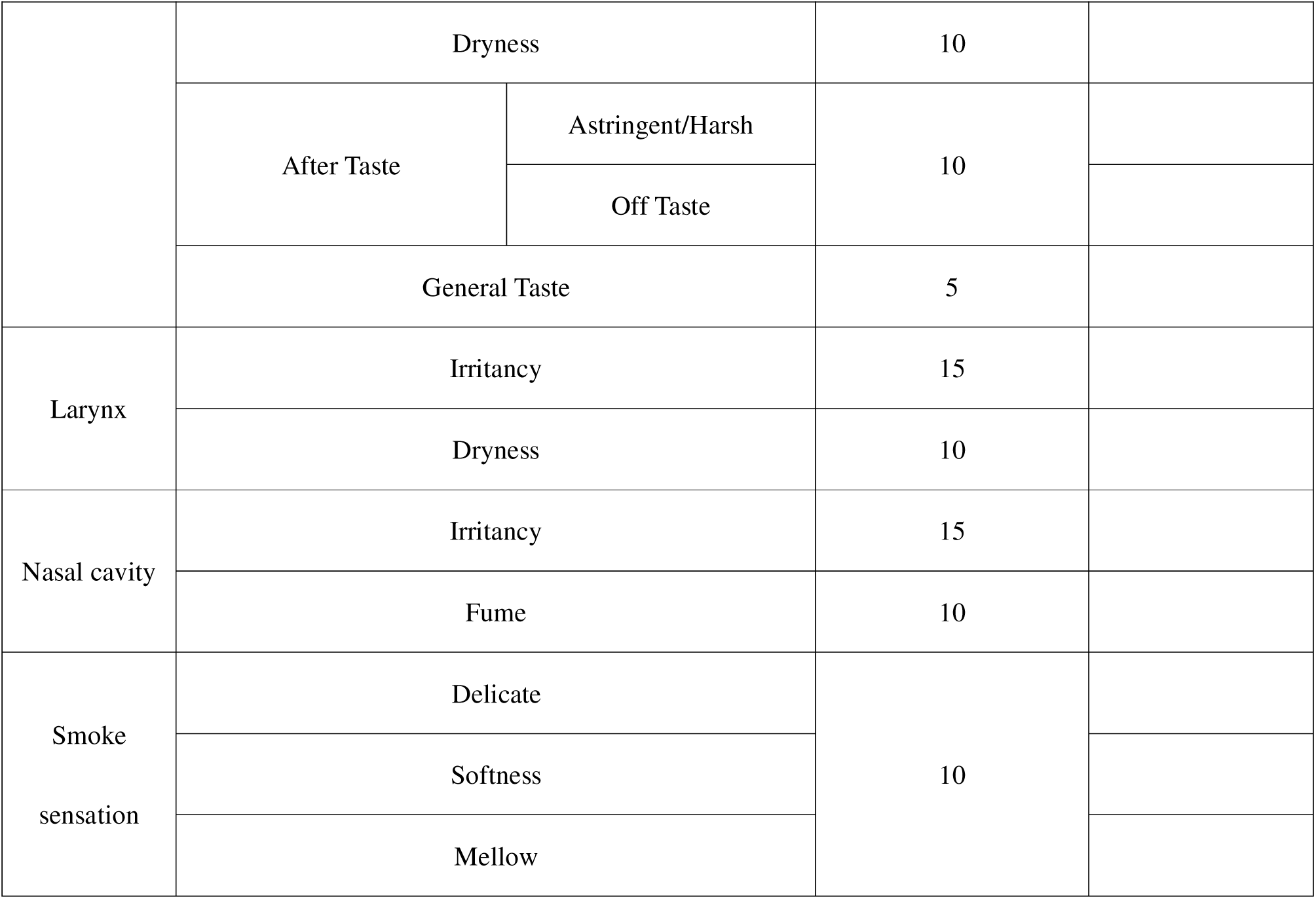
Evaluation method of cigarette smoking comfort.

## Results and Analysis

### Identification of Cigarettes Volatile Components Extracted by SAFE, SPME, and SDE

Volatile aroma compounds were isolated from Cambridge filters using three different extraction methods, SPME, SAFE and SDE, and the extracted volatile compounds were analyzed by gas chromatography-mass spectrometry (GC-MS). 89, 93 and 77 aroma compounds were detected by SPME, SAFE and SDE extraction methods, respectively.

**Table 2.** Identification of volatile compounds by SPME method.

| No. | Compound | CAS | c (mg/kg) |  |  |  |  |
| --- | --- | --- | --- | --- | --- | --- | --- |
|  |  |  | yk | ku | jy | hd | lj |
| 1 | 2-Methylpyridine | 109-06-8 | 8.6864±1.8312 | 9.3309±1.8954 | 9.0083±0.1055 | 3.6567±0.3672 | 6.8101±0.1635 |
| 2 | 2,6-Lutidine | 108-48-5 | 2.4245±0.1033 | 2.3271±0.0237 | 2.9899±0.0362 | 1.2567±0.0675 | 2.2674±0.1474 |
| 3 | 2-Ethylpyridine | 100-71-0 | 2.8876±0.0668 | 4.0810±0.8554 | 4.3527±1.2353 | 1.9567±0.1303 | 3.1311±0.5401 |
| 4 | 3-Methylpyridine | 108-99-6 | 20.8379±0.1484 | 23.4178±0.4595 | 26.6113±2.1887 | 15.567±1.7206 | 22.5581±1.9545 |
| 5 | 4-Methylpyridine | 108-89-4 | 3.2125±0.1401 | 3.5141±0.0303 | 4.0829±0.0345 | 2.3567±0.2779 | 3.3057±0.1373 |
| 6 | 2,5-Lutidine | 589-93-5 | 15.3234±5.4932 | 22.7543±1.0999 | 18.5887±6.6421 | 18.567±7.5831 | 11.8099±5.3482 |
| 7 | 2,4-Lutidine | 108-47-4 | 0 | 0 | 11.5843±0.0740 | 6.8567±0.7263 | 0 |
| 8 | Dimethyl malonate | 108-59-8 | 0 | 0 | 0 | 4.9567±0.1293 | 0 |
| 9 | 3-Ethylpyridine | 536-78-7 | 10.2149±0.2005 | 11.6235±0.0533 | 11.6861±0.1955 | 8.2567±1.2451 | 9.7482±0.5743 |
| 10 | 3,5-Dimethylpyridine | 591-22-0 | 7.7033±5.3612 | 14.1379±0.3799 | 0 | 2.1567±0.1783 | 5.0500±5.6294 |
| 11 | furan-2-carbaldehyde | 98-01-1 | 16.3446±1.8545 | 22.7147±2.4810 | 14.8990±8.0609 | 15.567±2.1431 | 18.2172±2.4923 |
| 12 | 3-Propylpyridine | 4673-31-8 | 6.5502±1.5632 | 6.9657±0.1734 | 5.8438±0.2577 | 7.4567±2.1543 | 4.8858±0.3588 |
| 13 | Ethyl nonanoate | 123-29-5 | 0 | 0 | 0 | 29.567±10.2667 | 0 |
| 14 | 3-methylcyclopent-2-en-1-one | 2758-18-1 | 16.9265±0.3268 | 31.9346±11.9094 | 22.3206±0.3091 | 18.567±2.0658 | 17.5516±1.0396 |
| 15 | 5-Methylfurfural | 620-02-0 | 29.8562±0.1289 | 46.7194±0.8381 | 39.8724±0.2318 | 35.567±3.7748 | 31.8528±1.4977 |
| 16 | 2-Methylbenzofuran | 4265-25-2 | 17.4057±8.2119 | 18.3673±1.6973 | 21.7801±8.9414 | 17.567±7.1626 | 13.8599±0.7519 |
| 17 | Furfuryl alcohol | 98-00-0 | 0 | 35.0528±0.0269 | 28.9415±0.0757 | 16.567±1.9334 | 15.5202±0.5781 |
| 18 | Crotonic acid | 3724-65-0 | 0 | 13.1580±0.4979 | 0 | 10.567±1.8739 | 0 |
| 19 | Naphthalene | 91-20-3 | 31.6064±0.6675 | 0 | 36.6698±1.3237 | 32.567±8.5084 | 30.2683±1.6325 |
| 20 | 2-Hydroxy-3,4-dimethyl-2-cyclopenten-1-one | 21835-00-7 | 7.6685±0.4436 | 0 | 9.5025±0.2408 | 14.567±2.2524 | 7.9942±0.5060 |
| 21 | 2-Hydroxy-3-methylcyclopent-2-enone | 80-71-7 | 34.0773±0.7999 | 54.7572±0.6587 | 39.7192±0.5759 | 50.567±5.1688 | 38.2581±1.6590 |
| 22 | 2-Methylnaphthalene | 91-57-6 | 33.3890±0.4031 | 34.0428±16.1977 | 45.2450±22.5189 | 42.567±4.9277 | 30.1418±15.5423 |
| 23 | α-terpinene | 99-86-5 | 0 | 0 | 0 | 12.567±5.1880 | 0 |
| 24 | Quinoline | 91-22-5 | 19.6857±1.4894 | 0 | 29.8056±15.4441 | 24.567±2.1052 | 0 |
| 25 | Maltol | 118-71-8 | 0 | 35.0711±0.5541 | 25.5885±0.9227 | 27.567±3.7870 | 20.2275±2.0358 |
| 26 | o-Cresol | 95-48-7 | 21.7940±0.4526 | 37.3492±0.3503 | 25.2297±0.5614 | 37.567±10.8459 | 33.5315±10.1314 |
| 27 | Phenol | 108-95-2 | 41.2466±0.7670 | 68.9959±0.9165 | 48.5564±0.7174 | 57.567±6.8447 | 45.2170±1.9728 |
| 28 | Isoquinoline | 119-65-3 | 0 | 0 | 0 | 22.567±3.3835 | 18.6539±2.0392 |
| 29 | 1,4-Dimethylnaphthalene | 571-58-4 | 0 | 0 | 0 | 24.567±3.1562 | 0 |
| 30 | Triacetin | 102-76-1 | 442.0674±6.9842 | 335.7248±13.9367 | 0 | 483567±37.3380 | 580.2923±20.5423 |
| 31 | 2,4-Dimethylphenol | 105-67-9 | 18.3109±0.5523 | 29.9636±0.2285 | 19.5805±0.6296 | 38.567±9.9540 | 25.4395±2.6688 |
| 32 | p-Cresol | 106-44-5 | 35.5503±19.1858 | 69.1481±21.7031 | 46.7914±14.7435 | 36.567±5.9024 | 39.6678±20.4821 |
| 33 | m-Cresol | 108-39-4 | 34.4036±0.9612 | 38.1714±0.7508 | 25.2192±1.6615 | 52.567±26.0204 | 0 |
| 34 | 1-methyl-1H-pyrrole-2-carbaldehyde | 1192-58-1 | 0 | 0 | 9.2554±1.3513 | 11.567±1.1823 | 8.6269±1.8643 |
| 35 | 4-Ethylphenol | 123-07-9 | 0 | 0 | 0 | 29.567±23.0688 | 0 |
| 36 | 2-Ethylphenol | 90-00-6 | 27.5867±14.0915 | 28.3777±17.1756 | 0 | 21.567±6.3374 | 0 |
| 37 | 2-Methoxy-4-vinylphenol | 7786-61-0 | 43.4400±3.1775 | 66.7206±1.0647 | 43.7259±3.2357 | 61.567±4.1963 | 46.9849±3.3176 |
| 38 | δ-elemene | 20307-84-0 | 0 | 0 | 12.7902±3.2458 | 14.567±2.8198 | 0 |
| 39 | 2,6-Dimethoxyphenol | 91-10-1 | 17.5536±1.3709 | 25.3622±0.8656 | 16.5047±1.7433 | 20.567±3.2876 | 16.8818±2.7779 |
| 40 | Benzyl benzoate | 120-51-4 | 0 | 0 | 0 | 49.567±38.3338 | 0 |
| 41 | R-Dihydroactinidiolide | 17092-92-1 | 5.2291±1.8542 | 9.6936±2.4170 | 4.6151±1.0528 | 5.7567±1.5815 | 5.8123±1.7407 |
| 42 | Indole | 120-72-9 | 6.2067±1.0863 | 15.0498±4.6011 | 0 | 15.567±1.0848 | 8.4277±1.0956 |
| 43 | 5-Hydroxymethylfurfural | 67-47-0 | 17.5963±1.9563 | 19.8166±1.1994 | 13.0661±1.5760 | 19.567±3.8652 | 21.8936±3.3643 |
| 44 | 2-Methylindole | 95-20-5 | 21.7320±2.8688 | 0 | 0 | 35.567±7.4636 | 0 |
| 45 | Palmitic acid | 57-10-3 | 44.5529±12.8206 | 60.5400±18.9240 | 43.0221±11.7426 | 78.567±11.0967 | 54.0399±10.0081 |
| 46 | Pyridine | 110-86-1 | 12.4644±0.2158 | 11.4662±0.0717 | 13.2007±0.0671 | 5.4567±0.3491 | 9.8272±0.4100 |
| 47 | Lemosol | 5989-27-5 | 17.2327±0.2406 | 13.0979±0.1310 | 14.5900±0.0966 | 3.8567±0.0697 | 11.2896±0.2130 |
| 48 | 1-Methylpyrrolidine | 120-94-5 | 0 | 2.8490±0.3034 | 0 | 3.7567±0.7626 | 0 |
| 49 | bornane | 464-15-3 | 0 | 0 | 0 | 6.2567±1.0112 | 0 |
| 50 | 3,4-Dimethyl-1,2-cyclopentanedione | 13494-06-9 | 0 | 0 | 0 | 23.567±0.2425 | 0 |
| 51 | 2,4,6-Trimethylphenol | 527-60-6 | 0 | 0 | 24.9802±11.5591 | 10.567±1.2642 | 0 |
| 52 | 2,3,6-Trimethylnaphthalene | 829-26-5 | 0 | 0 | 0 | 11.567±0.8689 | 0 |
| 53 | 3-Ethylphenol | 620-17-7 | 35.9955±19.5655 | 74.0613±21.6520 | 37.9831±21.6631 | 53.567±0.8005 | 54.1011±4.6278 |
| 54 | trans-Anethole | 4180-23-8 | 0 | 0 | 0 | 6.4567±0.2166 | 0 |
| 55 | 4-Vinylphenol | 2628-17-3 | 24.8113±2.6216 | 35.3059±1.8883 | 23.2548±3.0010 | 31.567±1.9979 | 25.1102±3.0052 |
| 56 | 4-Methyl-2-pentanone | 108-10-1 | 0.3723±0.0127 | 0 | 0 | 0 | 1.2051±1.0521 |
| 57 | β-Pinene | 18172-67-3 | 0.6673±0.0687 | 0 | 0 | 0 | 0 |
| 58 | Cyclopentanone | 120-92-3 | 1.1779±0.0066 | 0 | 0 | 0 | 0 |
| 59 | b-Ocimene | 13877-91-3 | 2.9755±0.6394 | 0 | 0 | 0 | 3.0700±0.0620 |
| 60 | 1,2,4-Trimethylbenzene | 95-63-6 | 2.8384±0.2955 | 12.2798±1.2895 | 5.1994±3.8496 | 0 | 7.6110±2.1116 |
| 61 | Non-8-en-2-one | 5009-32-5 | 2.9659±0.0323 | 0 | 0 | 0 | 0 |
| 62 | propanoic acid | 79-09-4 | 14.3285±5.1633 | 0 | 13.3412±1.1033 | 0 | 0 |
| 63 | 2-Methoxy-3-methylpyrazine | 2847-30-5 | 19.6751±0.0331 | 0 | 0 | 0 | 0 |
| 64 | Acetophenone | 98-86-2 | 7.3937±0.4082 | 0 | 0 | 0 | 0 |
| 65 | 1,2-Dihydro-1,1,6-trimethylnaphthalene | 30364-38-6 | 11.2112±0.2895 | 16.0483±0.2301 | 12.5057±1.0333 | 0 | 0 |
| 66 | Skatole | 83-34-1 | 25.3400±11.4988 | 32.6603±1.1768 | 28.5372±13.8046 | 0 | 35.3810±14.1350 |
| 67 | 2-Methoxy-4-methylphenol | 93-51-6 | 10.4700±0.5674 | 19.2886±1.7223 | 13.7667±1.3314 | 0 | 11.1270±0.0002 |
| 68 | 2,3-Xylenol | 526-75-0 | 24.3021±0.2423 | 33.5429±9.3426 | 0 | 0 | 0 |
| 69 | 1,6,7-Trimethylnaphthalene | 2245-38-7 | 7.9981±1.1835 | 0 | 0 | 0 | 0 |
| 70 | 4-Methoxy-1-ethylbenzene | 1515-95-3 | 10.4335±0.2900 | 0 | 0 | 0 | 0 |
| 71 | Benzoic acid | 65-85-0 | 17.5899±4.2925 | 25.5320±0.0776 | 0 | 0 | 18.7335±6.0760 |
| 72 | Styrene Monomer | 100-42-5 | 2.5058±0.0988 | 0 | 2.8593±0.0978 | 0 | 1.4526±0.0169 |
| 73 | 1,2,4,5-Tetramethyl benzene | 95-93-2 | 13.7280±0.4460 | 0 | 0 | 0 | 0 |
| 74 | 2-Methylbutanoic acid | 116-53-0 | 0 | 0 | 8.3722±6.9604 | 0 | 0 |
| 75 | 2-ethylfuran | 3208-16-0 | 0 | 0 | 3.9887±0.2269 | 0 | 0 |
| 76 | 2-Acetylfuran | 1192-62-7 | 0 | 0 | 16.4719±0.2596 | 0 | 12.8903±0.6631 |
| 77 | 2,6-Dimethylphenol | 576-26-1 | 0 | 0 | 23.2988±4.5834 | 0 | 0 |
| 78 | 2,5-Dimethylphenol | 95-87-4 | 0 | 0 | 13.2889±4.2138 | 0 | 0 |
| 79 | gamma-Terpineol | 586-81-2 | 0 | 0 | 8.5204±4.7662 | 0 | 0 |
| 80 | Isoeugenol | 5932-68-3 | 0 | 18.9759±0.1742 | 11.2542±2.3444 | 0 | 11.6156±2.8927 |
| 81 | 1,3,6-Octatriene,3,7-dimethyl-, (3Z)- | 3338-55-4 | 0 | 2.9826±1.0694 | 0 | 0 | 0 |
| 82 | 1-Methyl-4-prop-1-en-2-ylbenzene | 1195-32-0 | 0 | 10.4860±0.3247 | 0 | 0 | 0 |
| 83 | 1-Methylnaphthalene | 90-12-0 | 0 | 26.9404±1.0869 | 0 | 0 | 0 |
| 84 | 2,6-Dimethylnaphthalene | 581-42-0 | 0 | 21.8639±7.5555 | 0 | 0 | 0 |
| 85 | Cinnamonnitrile | 1885-38-7 | 0 | 41.8702±18.5500 | 0 | 0 | 18.1133±1.3055 |
| 86 | (E,Z)-2,6-dimethylocta-2,4,6-triene | 7216-56-0 | 0 | 42.5047±13.2567 | 0 | 0 | 0 |
| 87 | Thiophene | 110-02-1 | 0 | 0 | 0 | 0 | 1.8376±0.5376 |
| 88 | 4-Methylanisole | 104-93-8 | 0 | 0 | 0 | 0 | 9.7699±12.7507 |
| 89 | α-Thujone | 546-80-5 | 0 | 0 | 0 | 0 | 17.1802±0.0373 |

**Table 3.**
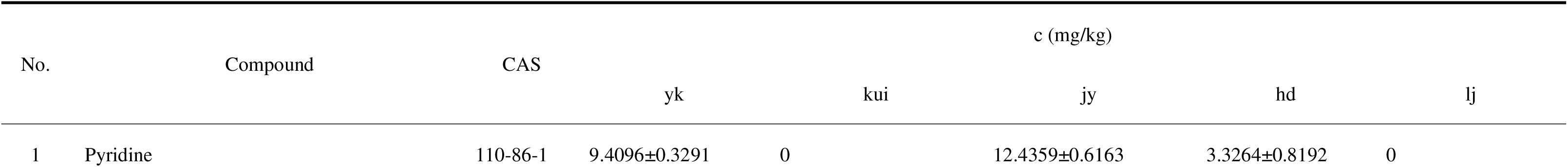
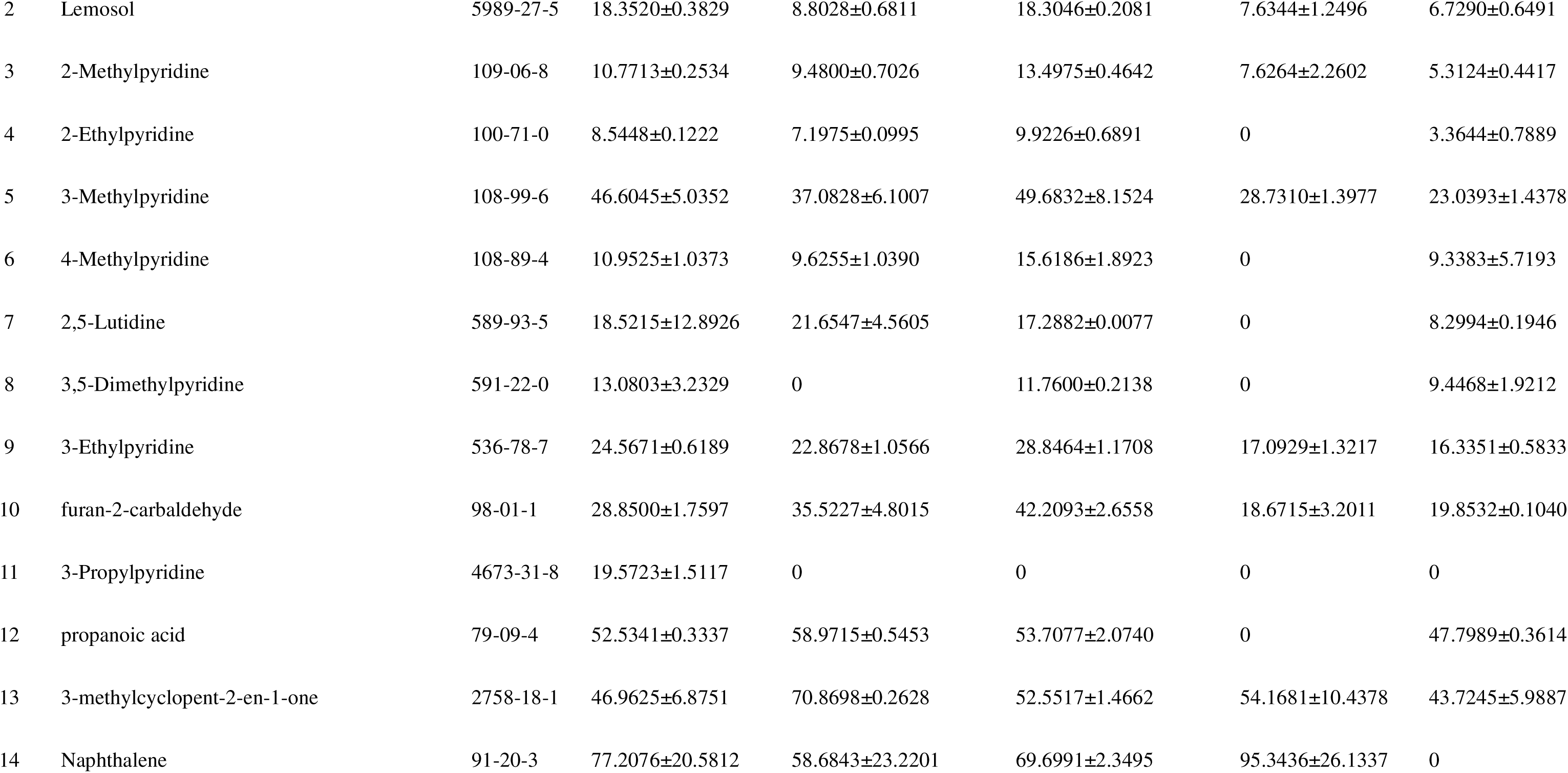
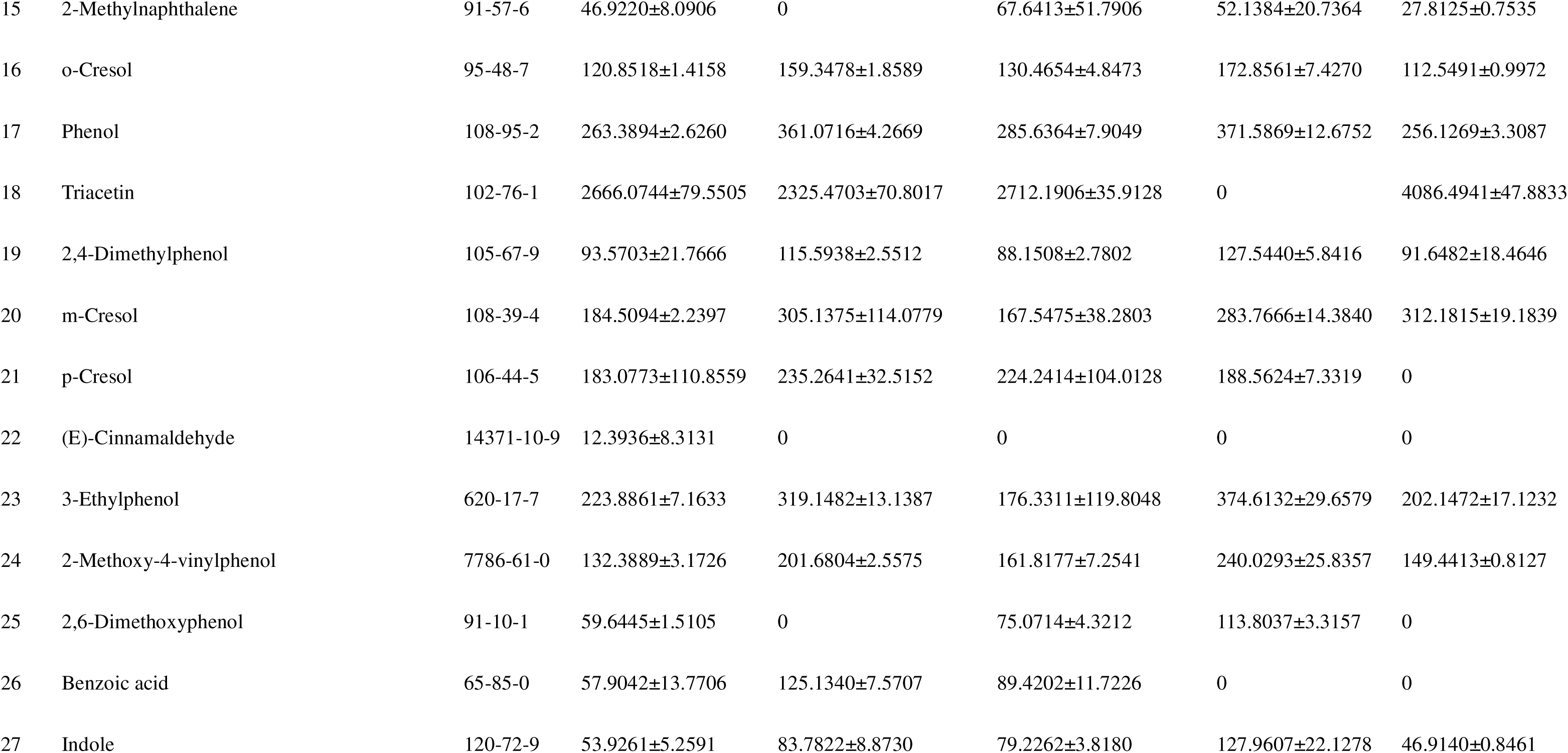
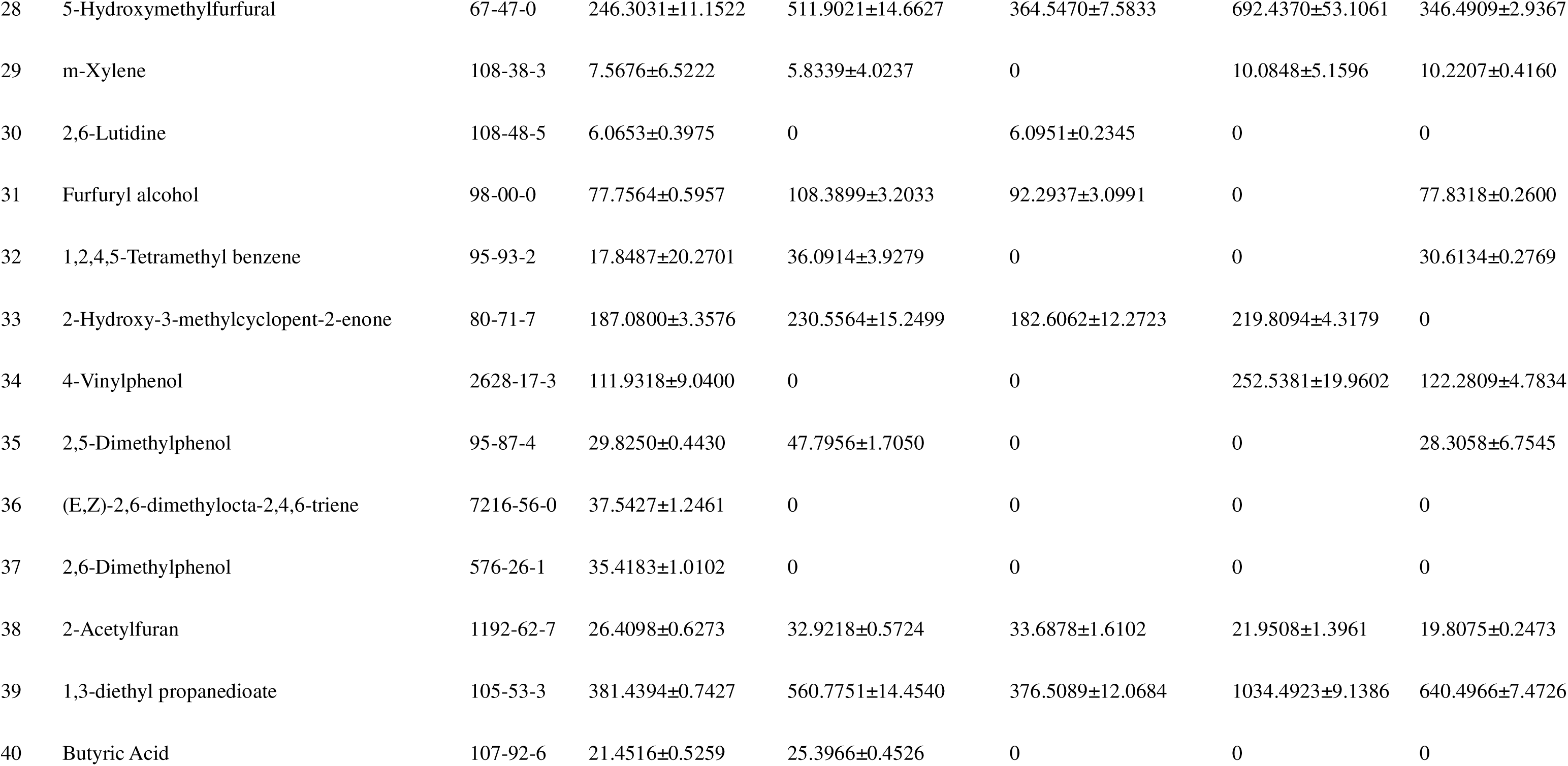
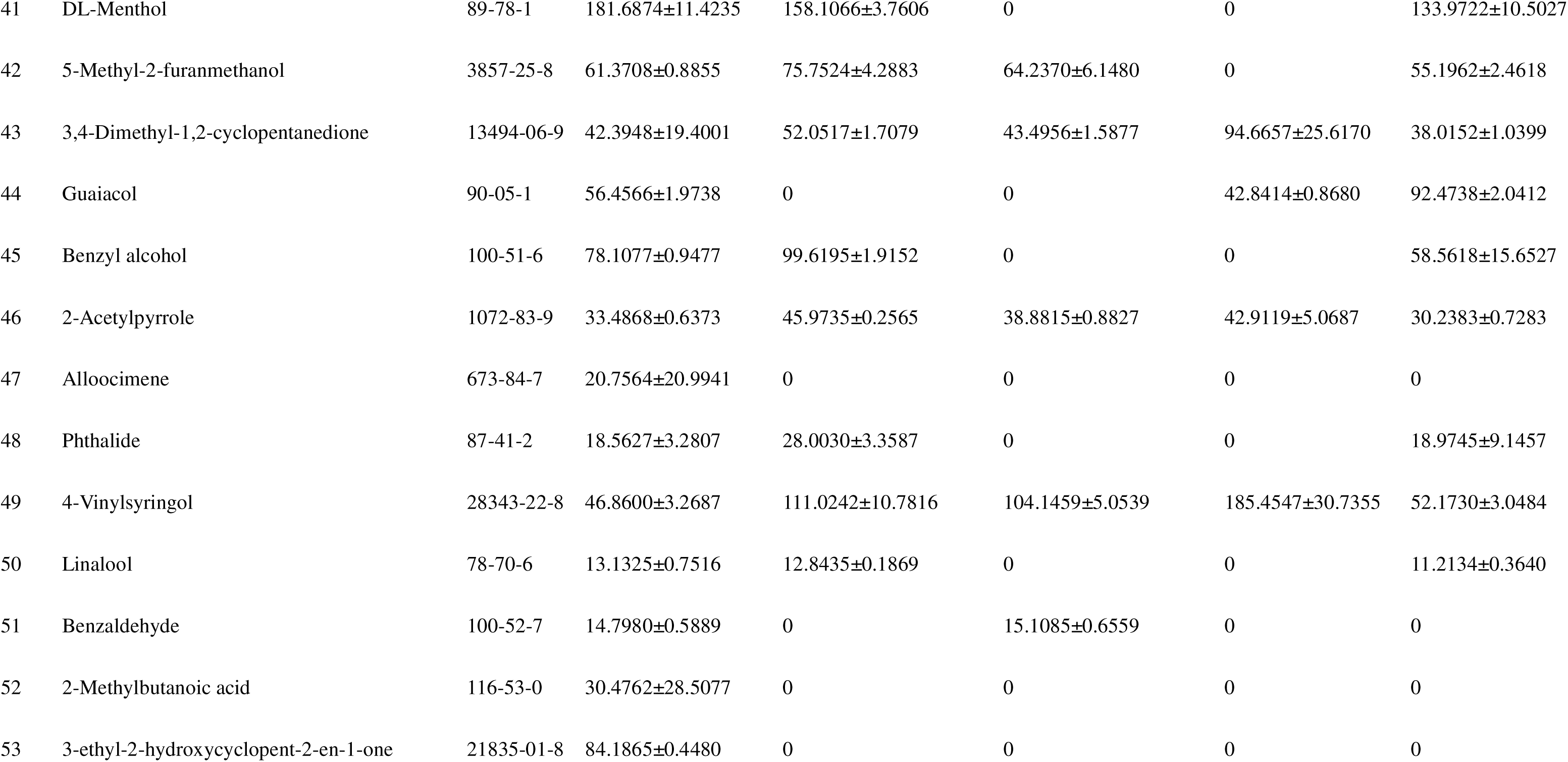
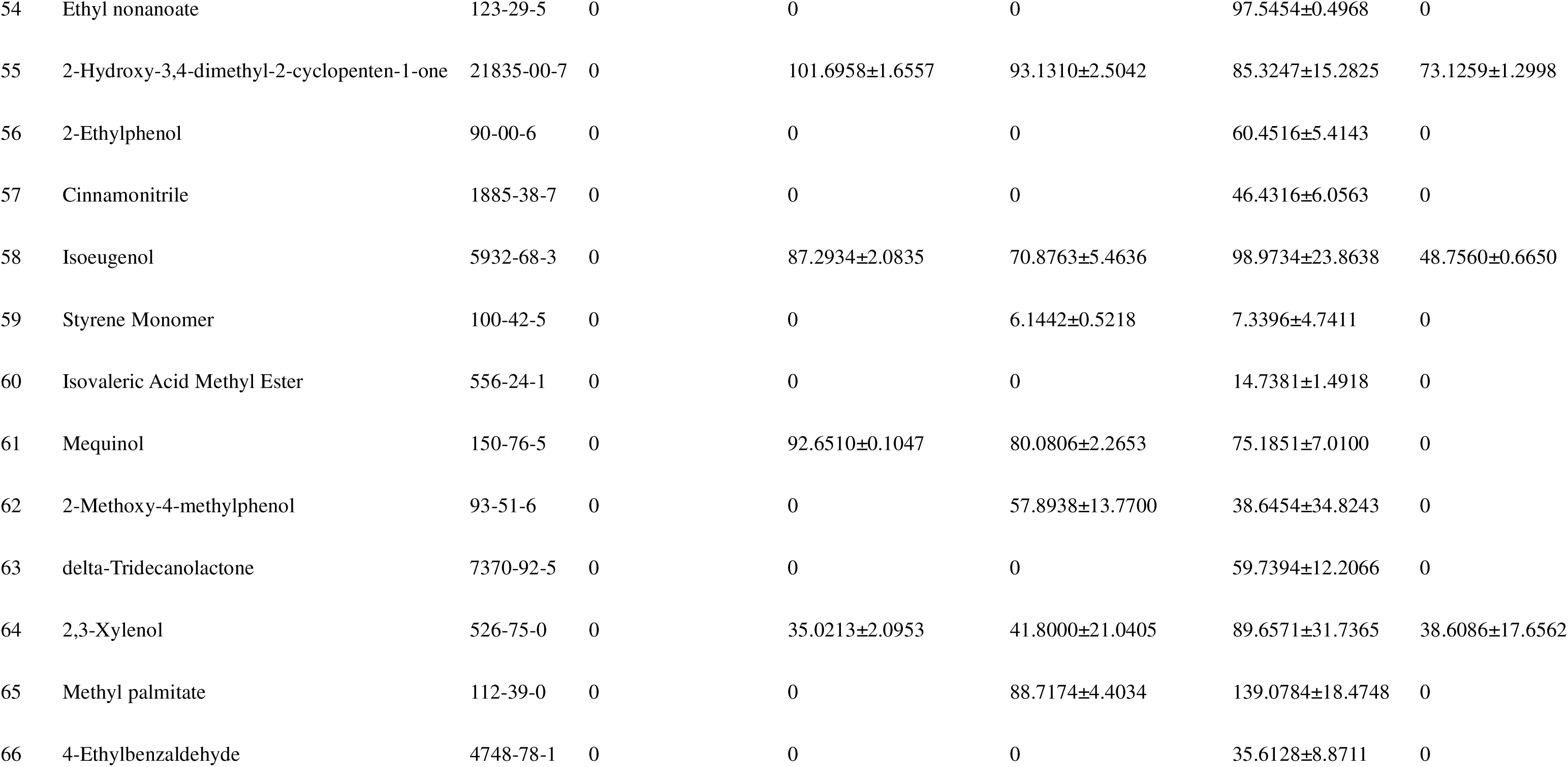
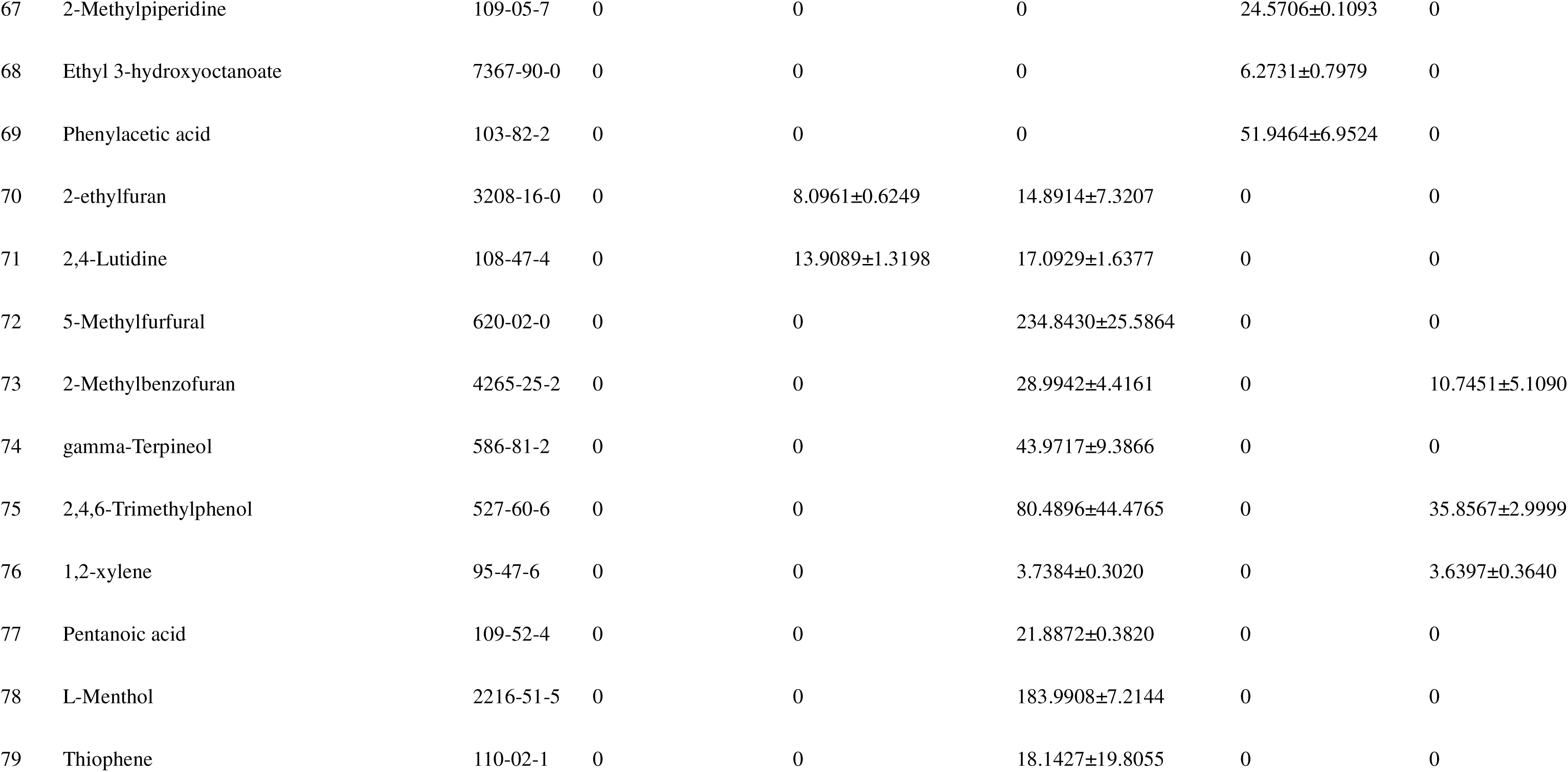
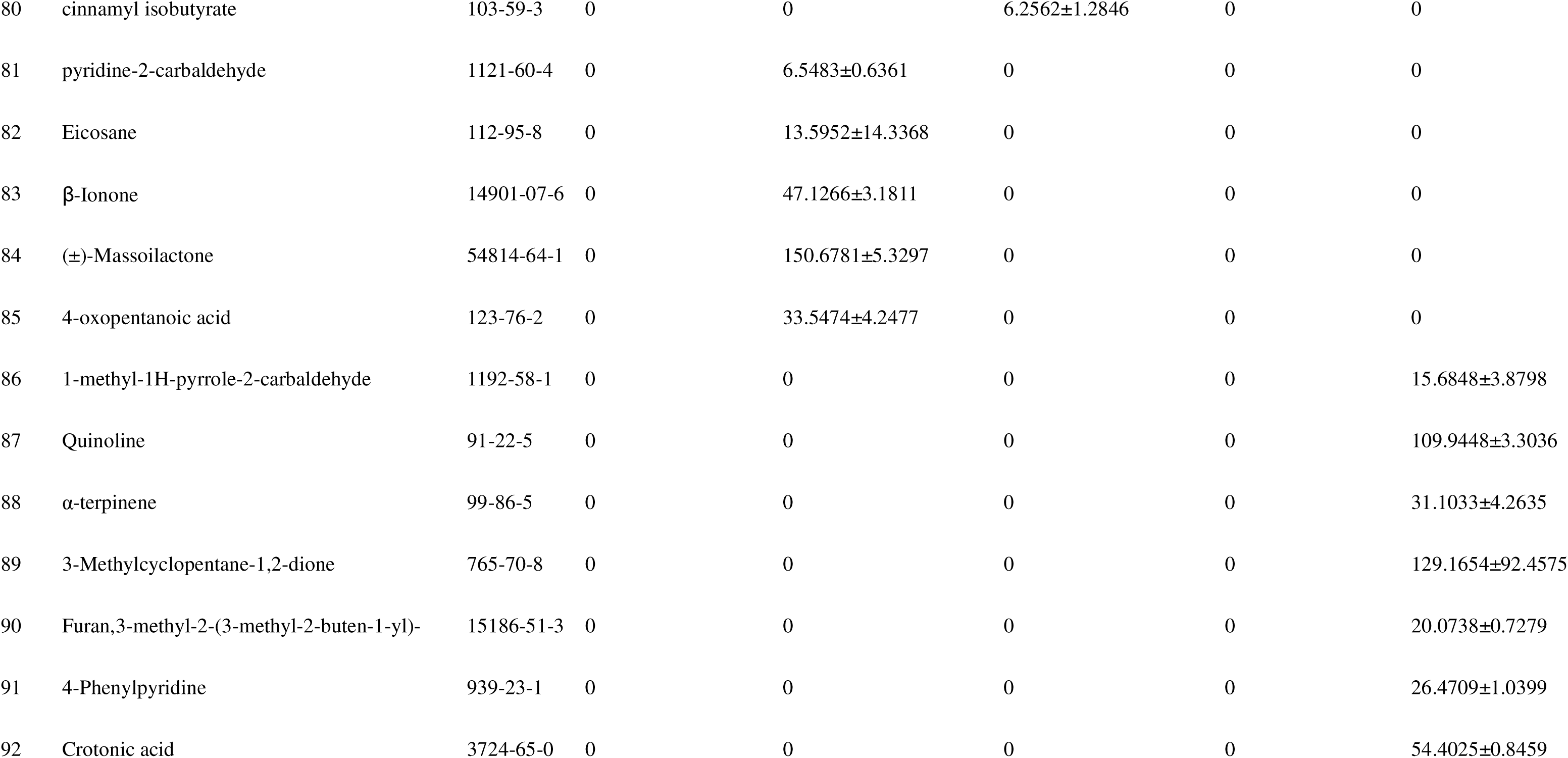
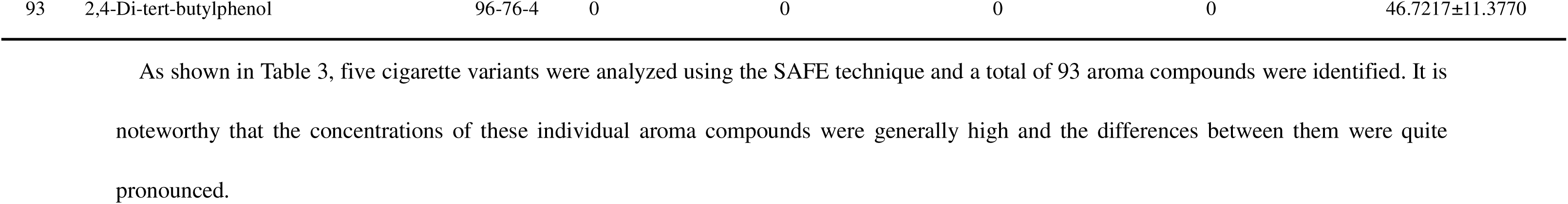
Identification of volatile compounds by SAFE method.

**Table 4.** Identification of volatile compounds by SDE method.

| No. | Compound | CAS | c (mg/kg) |  |  |  |  |
| --- | --- | --- | --- | --- | --- | --- | --- |
|  |  |  | yk | kui | jy | hd | lj |
| 1 | 1,2-xylene | 95-47-6 | 3.2929±0.3024 | 0 | 0 | 2.4781±0.0798 | 0 |
| 2 | Pyridine | 110-86-1 | 2.2166±0.0548 | 0 | 3.5669±0.0657 | 1.6424±0.0207 | 3.5669±0.0657 |
| 3 | Dodecane | 112-40-3 | 5.4098±1.3628 | 7.1866±2.0139 | 0 | 3.6130±0.0277 | 0 |
| 4 | Lemosol | 5989-27-5 | 2.8602±0.1958 | 3.3098±0.1209 | 3.4644±0.0698 | 2.9820±0.2076 | 3.4644±0.0698 |
| 5 | 2,3-Dimethylpyridine | 583-61-9 | 1.4832±0.2639 | 0 | 0 | 1.3901±0.6301 | 0 |
| 6 | Styrene Monomer | 100-42-5 | 1.5130±0.0388 | 2.3518±0.1451 | 0 | 0 | 0 |
| 7 | 2-Ethylpyridine | 100-71-0 | 2.2593±0.1114 | 0 | 2.4760±0.0095 | 1.0744±0.0085 | 2.4760±0.0095 |
| 8 | 3-Methylpyridine | 108-99-6 | 8.2863±1.6142 | 4.8283±0.9244 | 4.4774±5.3891 | 7.0415±0.2723 | 4.4774±5.3891 |
| 9 | 2,4-Lutidine | 108-47-4 | 1.8578±0.0475 | 1.5960±0.0734 | 0 | 0 | 0 |
| 10 | 2,5-Lutidine | 589-93-5 | 1.9551±0.3461 | 2.1396±1.2252 | 7.9174±7.3805 | 2.3406±0.0363 | 7.9174±7.3805 |
| 11 | n-Tetradecane | 629-59-4 | 8.1699±0.1139 | 13.0979±0.1682 | 10.8364±0.0495 | 9.6362±0.0043 | 10.8364±0.0495 |
| 12 | 2-Acetylfuran | 1192-62-7 | 3.9681±0.1046 | 0 | 0 | 3.9179±0.1282 | 0 |
| 13 | 3-methylcyclopent-2-en-1-one | 2758-18-1 | 10.6386±0.4186 | 7.7514±0.1948 | 28.1626±5.8276 | 10.0668±0.2398 | 28.1626±5.8276 |
| 14 | 5-Methylfurfural | 620-02-0 | 10.4745±0.4270 | 13.0845±0.4632 | 19.4224±0.6848 | 10.5018±0.2729 | 19.4224±0.6848 |
| 15 | Naphthalene | 91-20-3 | 7.3843±1.9052 | 10.1608±0.0288 | 8.1262±0.0597 | 6.6504±0.1812 | 8.1262±0.0597 |
| 16 | 2-Hydroxy-3,4-dimethyl-2-cyclopenten-1-one | 21835-00-7 | 6.7872±0.0770 | 11.6914±0.0350 | 13.6914±0.0448 | 0 | 13.6914±0.0448 |
| 17 | 2-Hydroxy-3-methylcyclopent-2-enone | 80-71-7 | 5.4512±0.4325 | 0 | 15.8393±0.0062 | 0 | 15.8393±0.0062 |
| 18 | Mequinol | 150-76-5 | 7.6847±6.9963 | 0 | 26.1585±0.1080 | 0 | 26.1585±0.1080 |
| 19 | Butylated hydroxytoluene | 128-37-0 | 43.1931±0.6091 | 78.1412±2.9347 | 57.6958±0.9939 | 48.3215±0.0415 | 57.6958±0.9939 |
| 20 | Quinoline | 91-22-5 | 8.8141±0.3315 | 0 | 0 | 0 | 0 |
| 21 | o-Cresol | 95-48-7 | 18.5087±0.3683 | 28.8234±0.2322 | 61.5630±6.3578 | 18.7132±0.4425 | 61.5630±6.3578 |
| 22 | Furan,3-methyl-2-(3-methyl-2-buten-1-yl)- | 15186-51-3 | 5.7195±0.8124 | 0 | 0 | 0 | 0 |
| 23 | Triacetin | 102-76-1 | 171.0426±1.3936 | 78.5189±0.5753 | 294.1084±3.1513 | 0 | 294.1084±3.1513 |
| 24 | 2,5-Dimethylphenol | 95-87-4 | 6.1592±0.2833 | 10.6673±0.1034 | 10.5645±0.3044 | 6.5377±0.3069 | 10.5645±0.3044 |
| 25 | 2,4-Dimethylphenol | 105-67-9 | 14.0654±0.3100 | 21.7960±0.3537 | 49.2248±13.8396 | 14.1723±0.5179 | 49.2248±13.8396 |
| 26 | m-Cresol | 108-39-4 | 46.5243±19.0169 | 43.5789±5.2012 | 76.0552±29.7828 | 58.9639±2.6030 | 76.0552±29.7828 |
| 27 | p-Cresol | 106-44-5 | 41.8981±24.7445 | 58.4464±25.3445 | 0 | 0 | 0 |
| 28 | 2,3-Xylenol | 526-75-0 | 11.9733±4.3469 | 19.4784±12.3391 | 16.6367±0.9118 | 10.2674±2.3766 | 16.6367±0.9118 |
| 29 | 3-Ethylphenol | 620-17-7 | 51.8941±2.2843 | 82.0107±0.6571 | 72.9786±9.5997 | 50.4888±8.1464 | 72.9786±9.5997 |
| 30 | 2-Methoxy-4-vinylphenol | 7786-61-0 | 20.3749±0.9929 | 30.9637±0.0929 | 29.2980±0.1844 | 21.6336±0.8120 | 29.2980±0.1844 |
| 31 | 2,4-Di-tert-butylphenol | 96-76-4 | 27.9735±1.3448 | 53.7890±0.1060 | 45.5104±0.6012 | 35.6837±1.7324 | 45.5104±0.6012 |
| 32 | Isoeugenol | 5932-68-3 | 18.2685±2.6201 | 0 | 0 | 23.1081±3.0964 | 0 |
| 33 | R-Dihydroactinidiolide | 17092-92-1 | 15.8677±0.6701 | 0 | 17.5993±10.5653 | 19.9764±1.1955 | 17.5993±10.5653 |
| 34 | Indole | 120-72-9 | 13.8067±0.2938 | 36.5432±1.8734 | 52.9299±3.4778 | 25.2757±1.2785 | 52.9299±3.4778 |
| 35 | 4-Ethylbenzaldehyde | 4748-78-1 | 10.5052±1.9428 | 0 | 0 | 0 | 0 |
| 36 | Palmitic acid | 57-10-3 | 95.8221±4.6621 | 132.6231±1.8058 | 122.4762±7.4275 | 126.7737±12.5595 | 122.4762±7.4275 |
| 37 | 2-methyl-2-buten-1-ol | 4675-87-0 | 2.2983±0.1204 | 0 | 0 | 0 | 0 |
| 38 | 2-Methylpyridine | 109-06-8 | 3.0751±0.0311 | 1.6191±0.0645 | 3.7854±0.0777 | 2.6680±0.0755 | 3.7854±0.0777 |
| 39 | furan-2-carbaldehyde | 98-01-1 | 6.5409±0.0262 | 7.4474±0.0089 | 8.7314±0.0783 | 6.3153±0.1136 | 8.7314±0.0783 |
| 40 | Furfuryl alcohol | 98-00-0 | 11.2503±0.0348 | 0 | 0 | 11.6893±0.0888 | 0 |
| 41 | 3-ethyl-2-hydroxycyclopent-2-en-1-one | 21835-01-8 | 7.2200±0.1234 | 0 | 14.4362±5.3487 | 11.4618±5.6155 | 14.4362±5.3487 |
| 42 | (E/Z)-Geranylacetone | 689-67-8 | 5.8040±0.2892 | 13.0916±0.2541 | 0 | 0 | 0 |
| 43 | Guaiacol | 90-05-1 | 16.7442±0.0361 | 28.9198±0.8075 | 0 | 19.0982±1.3587 | 0 |
| 44 | 2-Methoxy-4-methylphenol | 93-51-6 | 5.3115±4.4234 | 0 | 9.7478±0.3603 | 8.8155±0.3464 | 9.7478±0.3603 |
| 45 | Phenol | 108-95-2 | 29.1083±0.0509 | 0 | 64.7678±2.9180 | 14.3487±18.4124 | 64.7678±2.9180 |
| 46 | (+)-Aromadendrene | 489-39-4 | 12.8476±6.2531 | 0 | 0 | 0 | 0 |
| 47 | 2-methyl-5-Benzofuranol | 6769-56-8 | 6.6967±5.5865 | 0 | 0 | 0 | 0 |
| 48 | Mesityl Oxide | 141-79-7 | 0 | 3.3098±0.2779 | 0 | 0 | 0 |
| 49 | 2,6-Lutidine | 108-48-5 | 0 | 0.9502±0.0399 | 0 | 1.2939±0.2687 | 0 |
| 50 | Eicosane | 112-95-8 | 0 | 6.4810±6.0131 | 0 | 3.5077±0.3613 | 0 |
| 51 | Nonanal | 124-19-6 | 0 | 2.3155±0.0196 | 0 | 0 | 0 |
| 52 | Benzaldehyde | 100-52-7 | 0 | 4.7078±0.2557 | 0 | 0 | 0 |
| 53 | Cinnamonnitrile | 1885-38-7 | 0 | 15.0481±0.0143 | 16.1185±0.3736 | 0 | 16.1185±0.3736 |
| 54 | (-)-Alloaromadendrene | 25246-27-9 | 0 | 24.4955±8.6446 | 21.8537±18.1733 | 14.5841±9.4095 | 21.8537±18.1733 |
| 55 | 4-Vinylphenol | 2628-17-3 | 0 | 48.7768±0.0953 | 0 | 0 | 0 |
| 56 | 2-Methylindole | 95-20-5 | 0 | 57.1227±0.5092 | 0 | 0 | 0 |
| 57 | 4-Methylpyridine | 108-89-4 | 0 | 1.0770±0.1573 | 8.3726±6.1897 | 1.0810±0.1521 | 8.3726±6.1897 |
| 58 | α-Thujone | 546-80-5 | 0 | 10.3507±0.0278 | 0 | 0 | 0 |
| 59 | 2,4,6-Trimethylphenol | 527-60-6 | 0 | 27.2894±20.2797 | 0 | 0 | 0 |
| 60 | Isoeugenol | 97-54-1 | 0 | 29.5315±3.6667 | 24.0680±0.5585 | 0 | 24.0680±0.5585 |
| 61 | 2-ethylfuran | 3208-16-0 | 0 | 0 | 2.7561±0.0156 | 1.0594±0.8435 | 2.7561±0.0156 |
| 62 | cyclohex-2-en-1-one | 930-68-7 | 0 | 0 | 5.7289±0.1152 | 0 | 5.7289±0.1152 |
| 63 | Acetophenone | 98-86-2 | 0 | 0 | 7.5564±0.0836 | 0 | 7.5564±0.0836 |
| 64 | 5-Methyl-2-furanmethanol | 3857-25-8 | 0 | 0 | 4.7694±0.2431 | 0 | 4.7694±0.2431 |
| 65 | 3-Carene | 13466-78-9 | 0 | 0 | 6.9451±0.2726 | 0 | 6.9451±0.2726 |
| 66 | 4-Methoxy-1-ethylbenzene | 1515-95-3 | 0 | 0 | 28.4775±0.2644 | 0 | 28.4775±0.2644 |
| 67 | (E,Z)-2,6-dimethylocta-2,4,6-triene | 7216-56-0 | 0 | 0 | 13.8857±2.8044 | 12.1273±2.0230 | 13.8857±2.8044 |
| 68 | Non-8-en-2-one | 5009-32-5 | 0 | 0 | 0.3721±0.3134 | 0 | 0.3721±0.3134 |
| 69 | Octadecanaldehyde | 638-66-4 | 0 | 0 | 5.7316±2.2559 | 0 | 5.7316±2.2559 |
| 70 | 2-Acetylpyrrole | 1072-83-9 | 0 | 0 | 18.8630±0.0377 | 11.9708±0.5206 | 18.8630±0.0377 |
| 71 | 2,6-Dimethylpyrazine | 108-50-9 | 0 | 0 | 0 | 0.4029±0.0197 | 0 |
| 72 | 2-Vinylpyridine | 100-69-6 | 0 | 0 | 0 | 7.7525±0.0831 | 0 |
| 73 | 2-Methoxy-3-methylpyrazine | 2847-30-5 | 0 | 0 | 0 | 10.9877±0.2454 | 0 |
| 74 | Isoquinoline | 119-65-3 | 0 | 0 | 0 | 7.2813±2.3517 | 0 |
| 75 | Phenyl acetate | 122-79-2 | 0 | 0 | 0 | 27.1579±0.8107 | 0 |
| 76 | 2-methoxy-5-prop-2-enyl-phenol | 501-19-9 | 0 | 0 | 0 | 14.2412±1.0300 | 0 |
| 77 | Skatole | 83-34-1 | 0 | 0 | 0 | 37.8050±3.1698 | 0 |

### Aroma components of five kinds of cigarettes were analyzed by GC×GC-MS

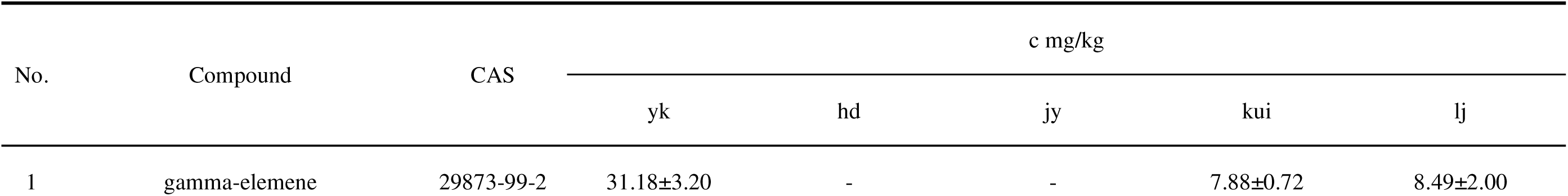
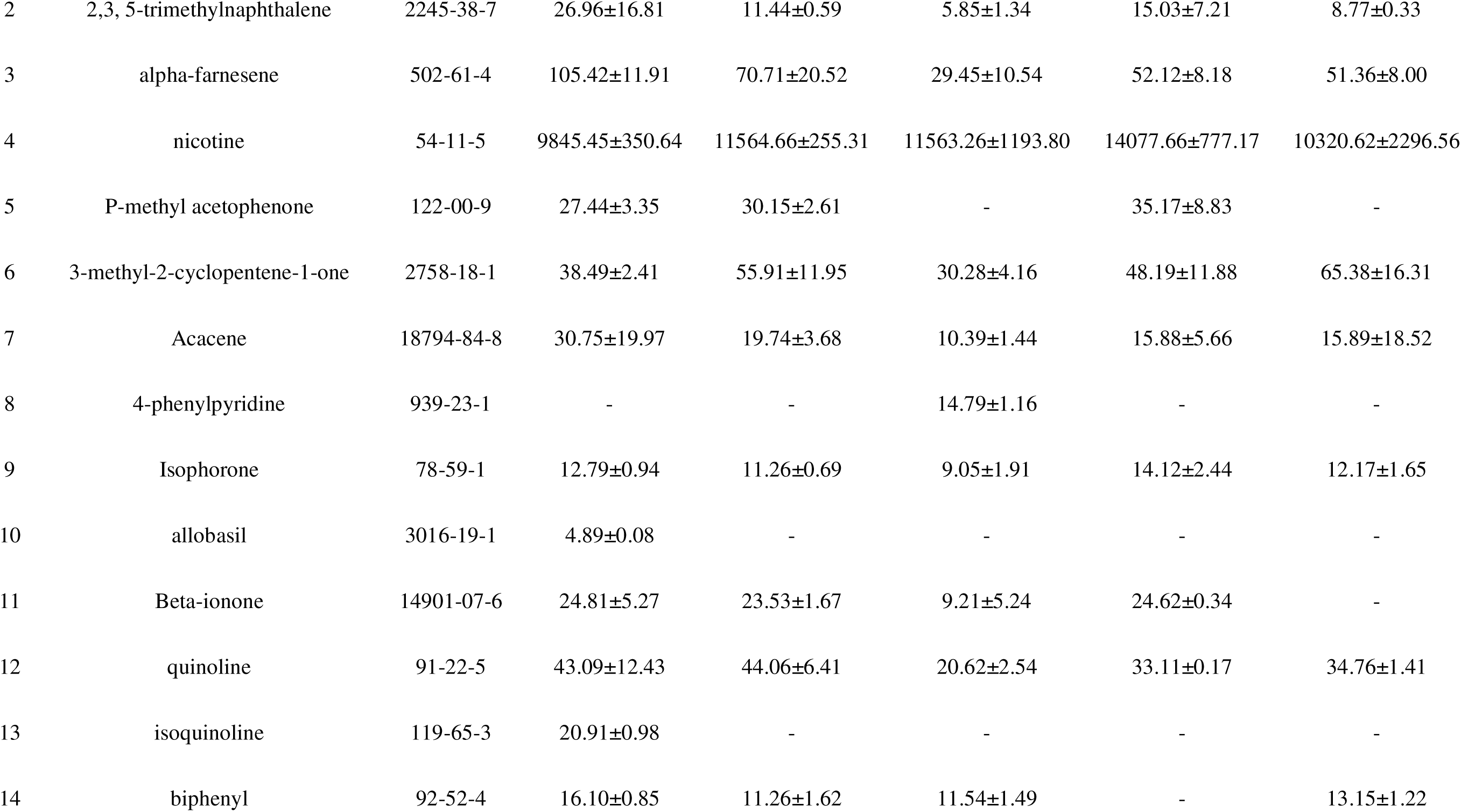
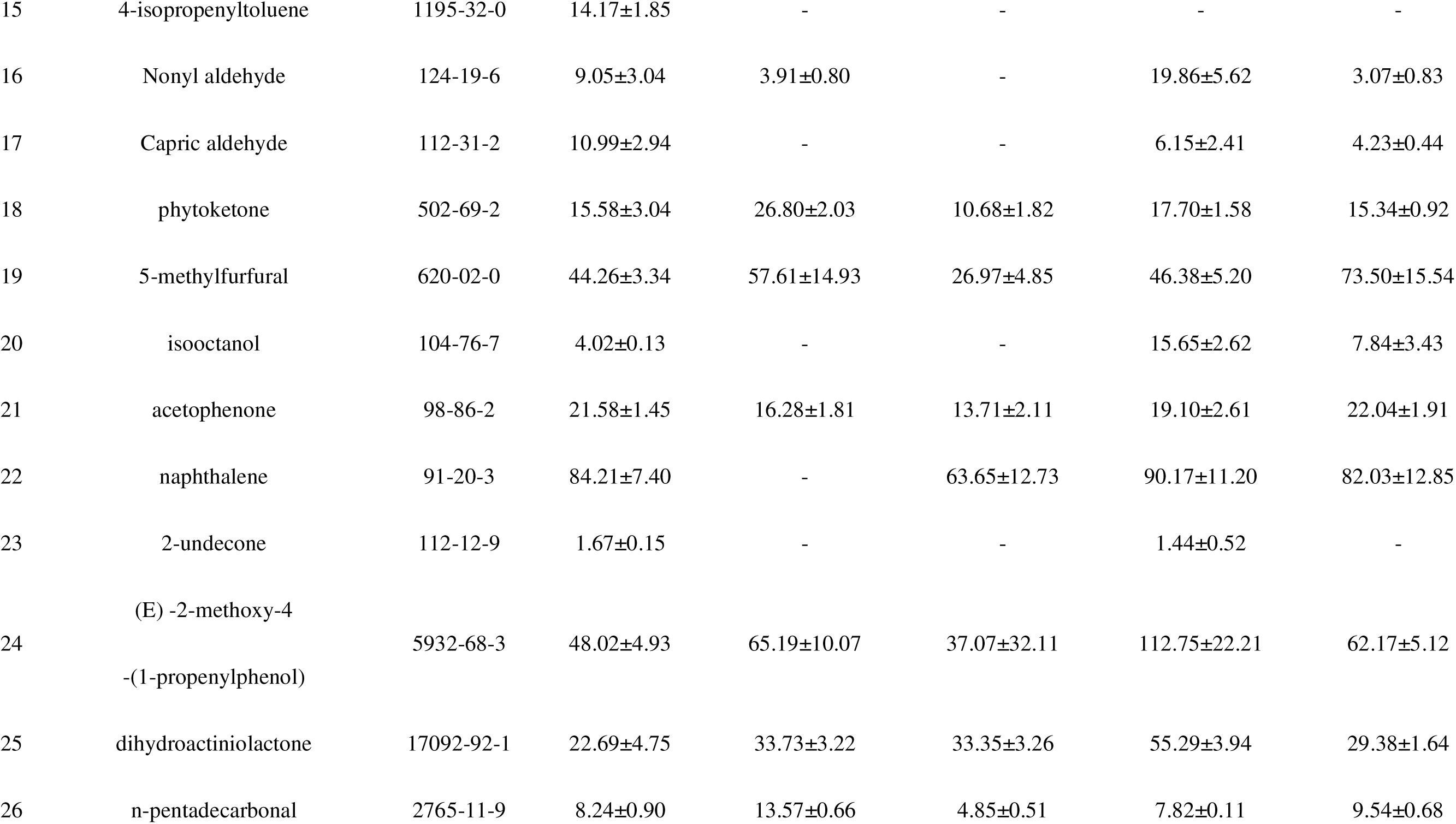
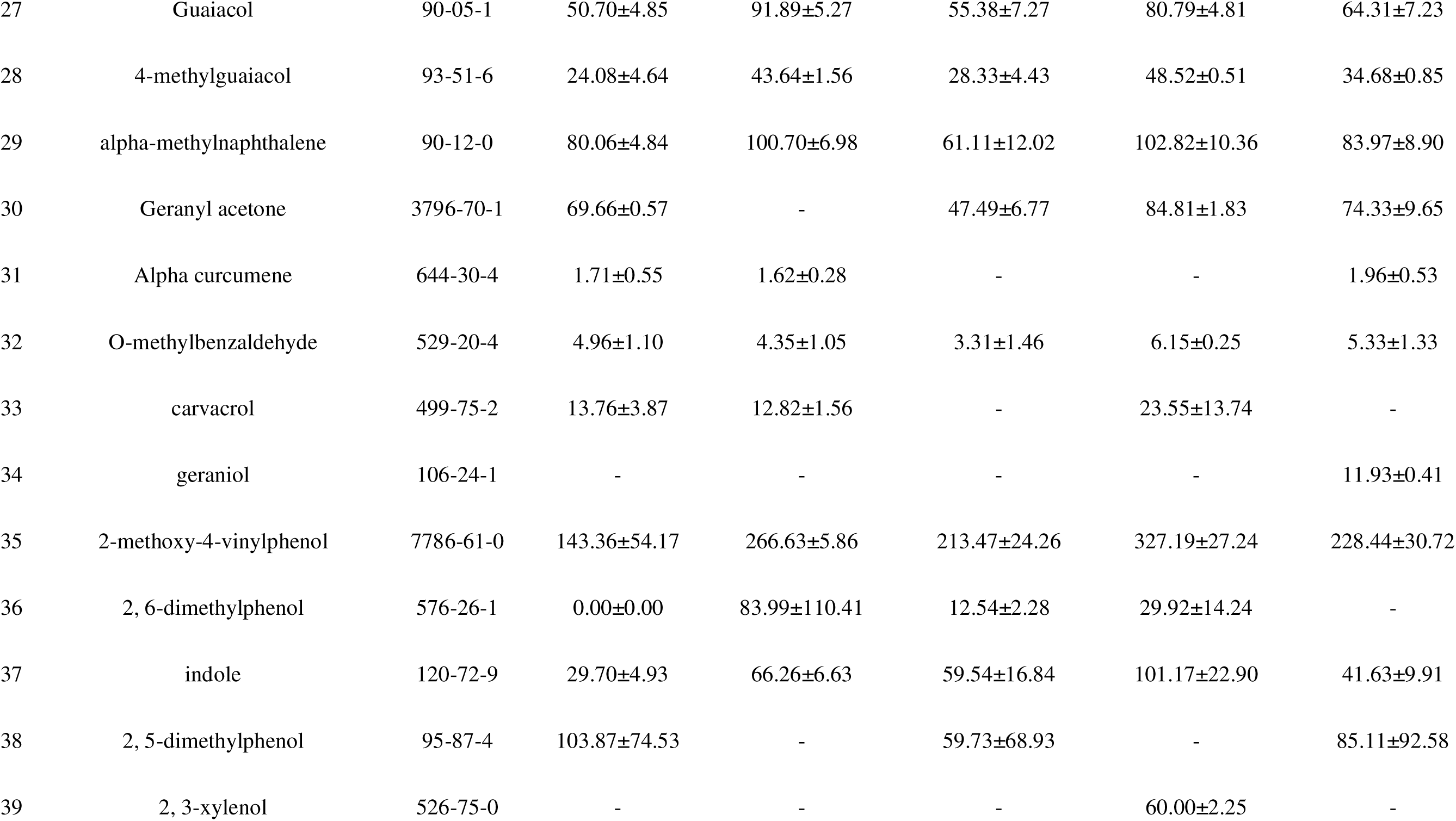
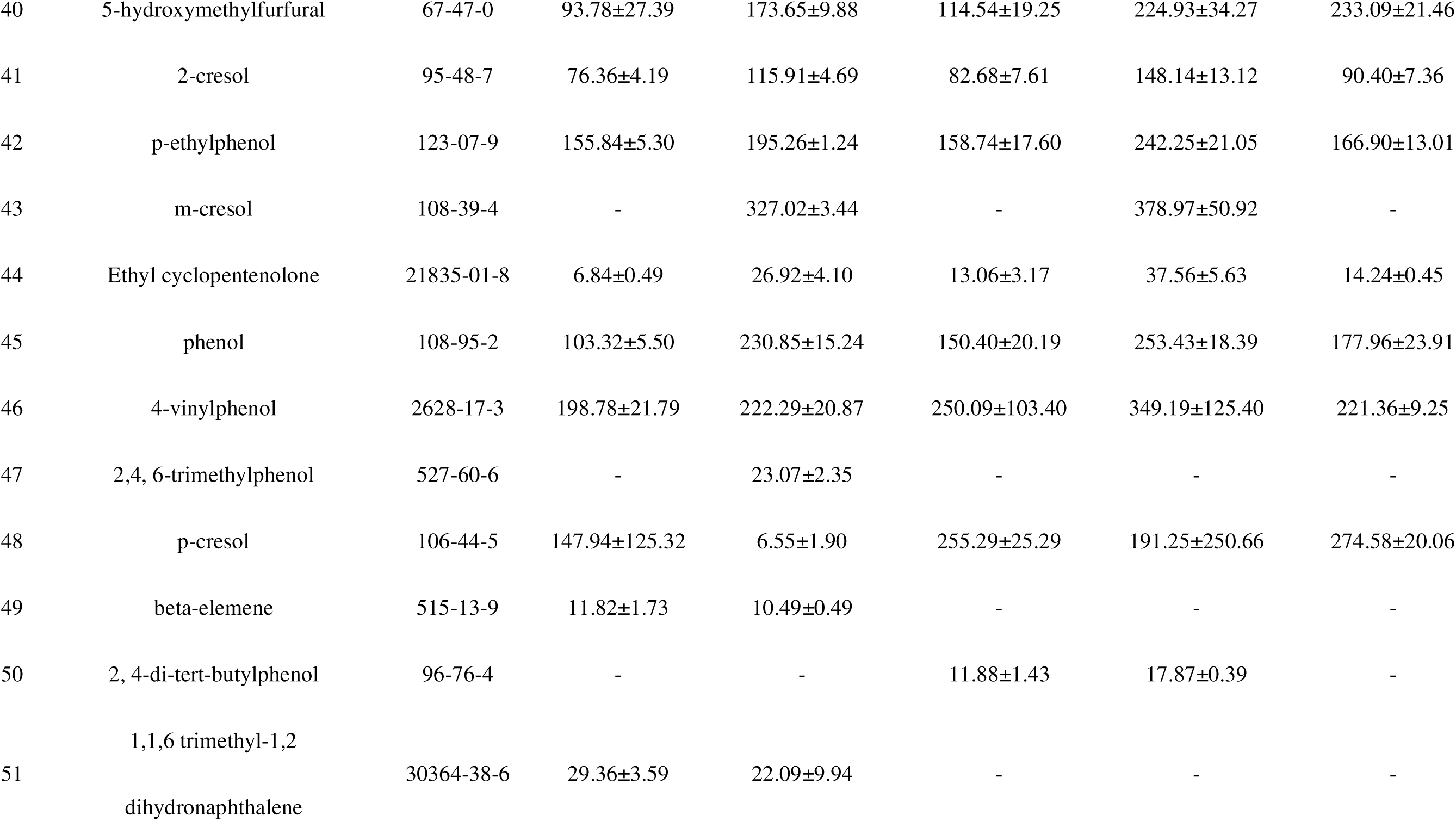
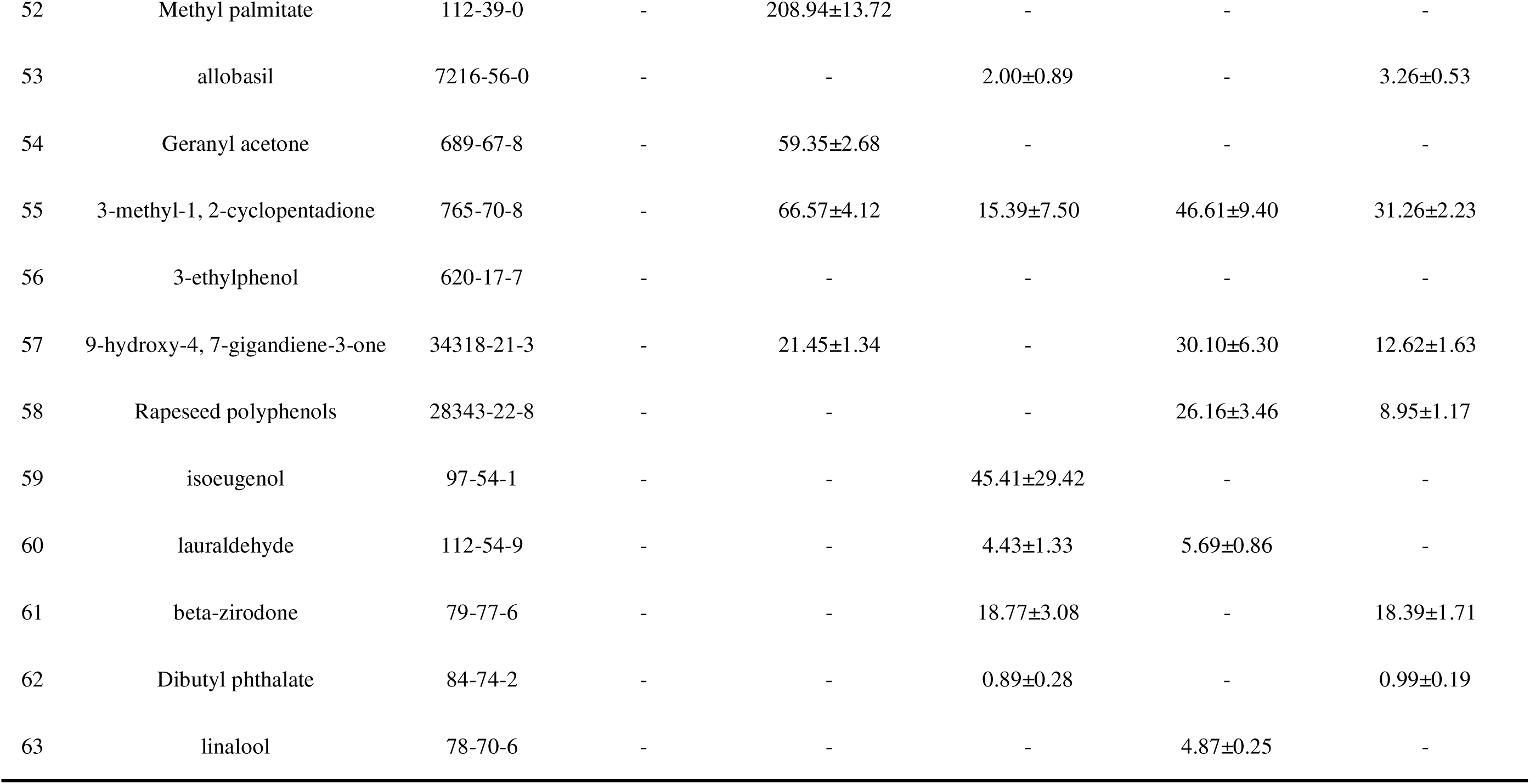

### Selection of pre-treatment methods

In this study, five different types of cigarette filters were extracted using three different pre-treatments (SAFE, SPME and SDE) and the volatile compounds in the extracts were analyzed using GC-MS. A total of 15 samples were performed and three parallel experiments were conducted for each sample. The content data obtained were compared and analyzed by principal component analysis (PCA).

**Figure 1.**
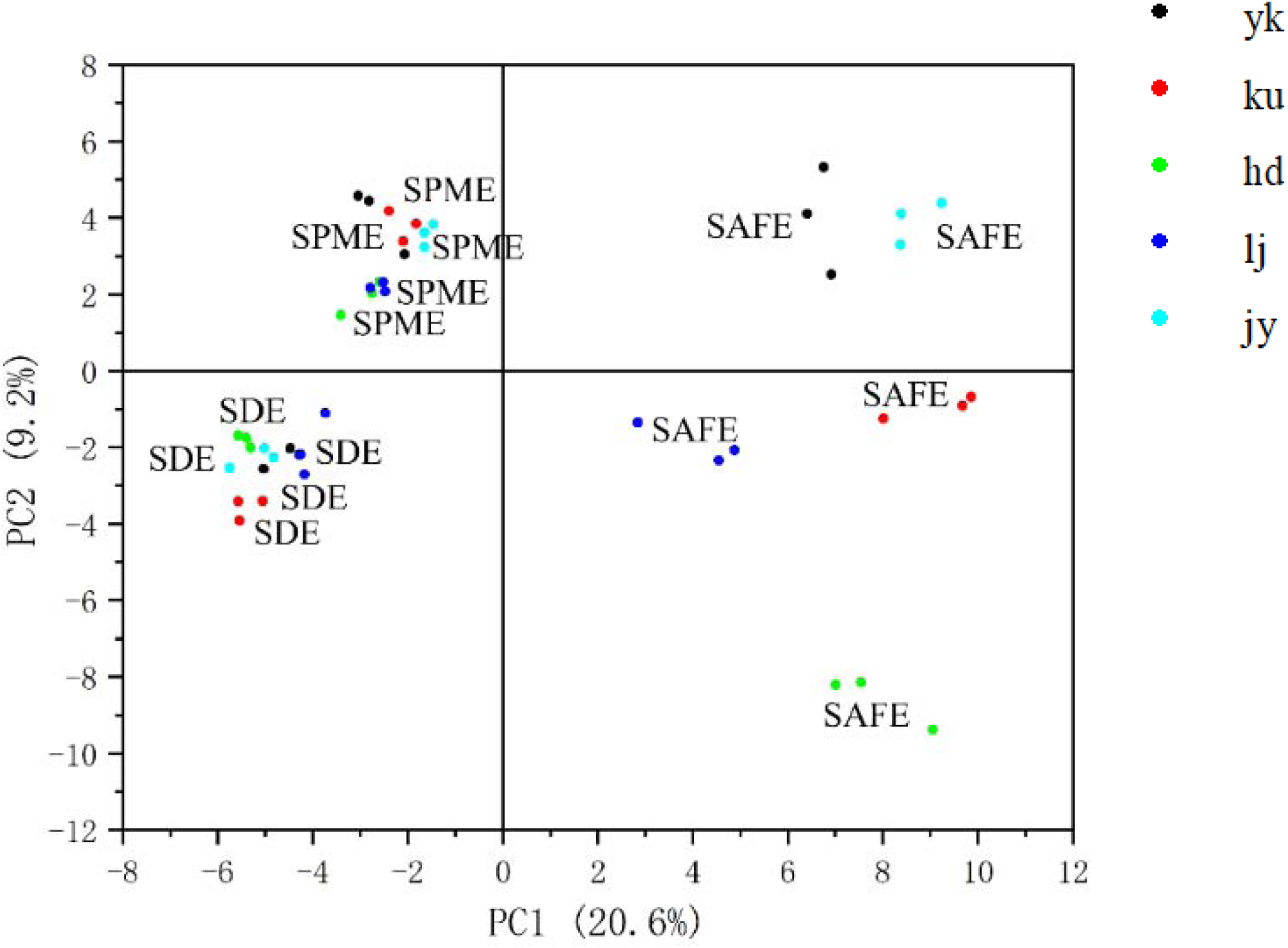
PCA analysis of the aroma components of five types of cigarettes by different pretreatment methods

GC-MS analysis was performed to compare the effectiveness of the three pretreatment methods in extracting volatile compounds from different types of cigarette filters. The PCA results showed that the SPME and SDE methods were not able to distinguish the aroma components of the five types of cigarettes very well, compared with the solvent-assisted flavor evaporation (SAFE) method, which showed a better differentiation ability. The PCA results clearly showed the aroma characteristics of each type of cigarettes. characteristics, which suggests that the method is more effective in extracting volatile compounds from cigarette filters. Therefore, we can conclude that solvent-assisted flavor evaporation extraction is a better pretreatment method, which is especially suitable for distinguishing the aroma substances of different types of cigarettes.

The aroma profiles of cigarettes are inherently complex, consisting of a wide range of volatile compounds with varying concentrations and interactions. This complexity can lead to a more dispersed distribution of variance across multiple principal components. While the cumulative variance explained by PC1 and PC2 is not exceptionally high, it still captures a significant portion of the variability in the dataset. The remaining variance is likely distributed across higher-order components, reflecting the intricate nature of the aroma compounds and their interactions.

These results provide important guidance for choosing a suitable pretreatment method, and also provide useful information for further understanding the extraction mechanism of volatile compounds in cigarettes.

**Figure 2.**
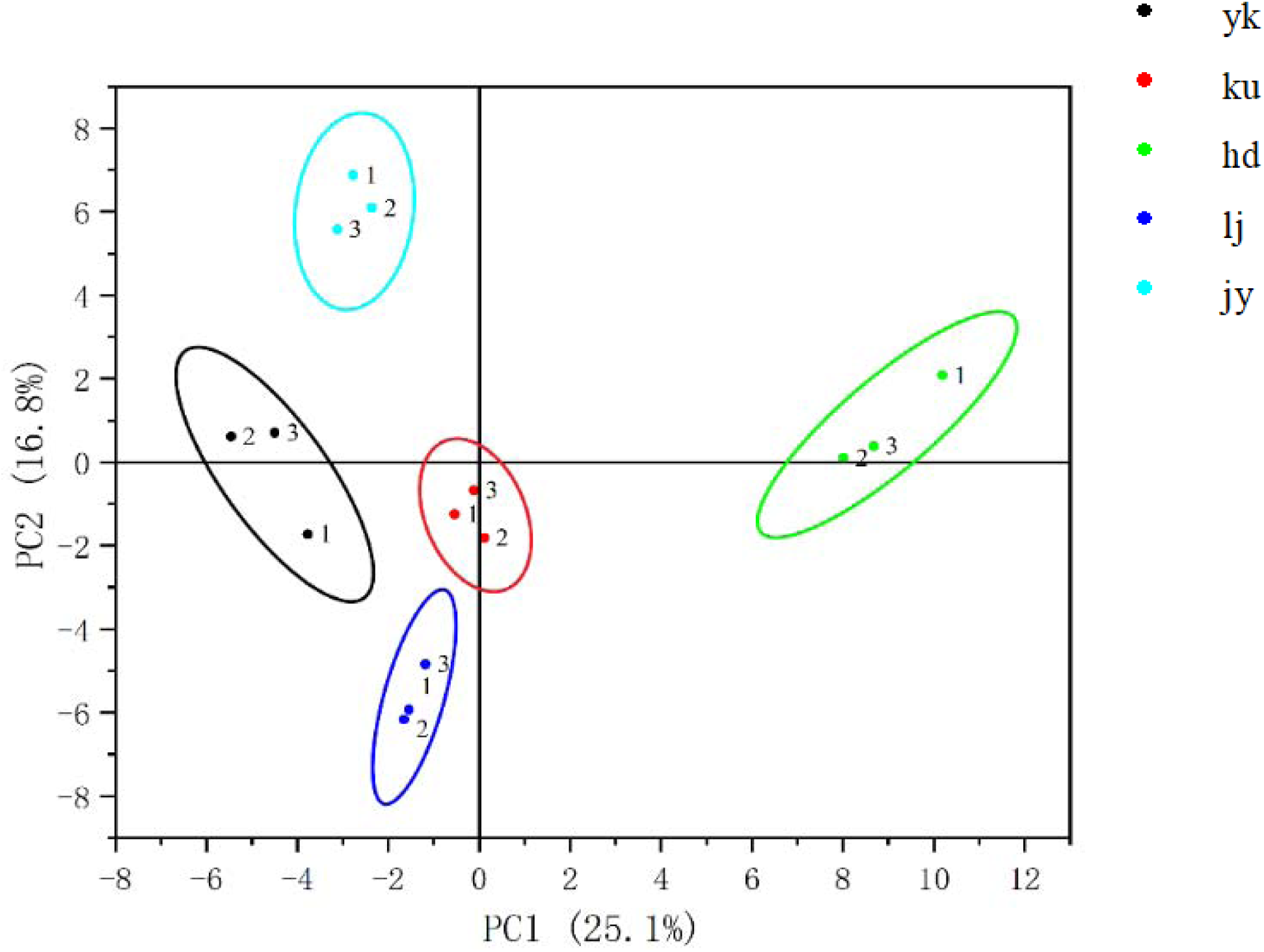
PCA analysis of the aroma components of five cigarettes by the SAFE pretreatment method

Through PCA analysis, we observed that the PCA results using the solvent-assisted flavor evaporation (SAFE) method clearly demonstrated significant differences between the aroma compositions of five different types of cigarettes. The data points corresponding to each type of cigarette sample were clustered in different regions, which indicates the unique characteristics of their volatile compound compositions. Therefore, the results of PCA analysis further validated the superiority of the solvent-assisted flavor evaporation method in extracting cigarette volatile compounds, and demonstrated that it is a reliable pretreatment method suitable for studying and differentiating the aroma profiles of different types of cigarettes.

### Differences in aroma of different cigarettes

By calculating the Odor Activity Values (OAVs) of aroma compounds extracted using the Solvent Assisted Flavor Evaporation (SAFE) method, we can identify compounds contributing to the aroma of cigarettes. Specifically, compounds with OAVs greater than or equal to 1 are retained as they are considered to significantly contribute to the aroma of cigarettes.

**Table 5.**
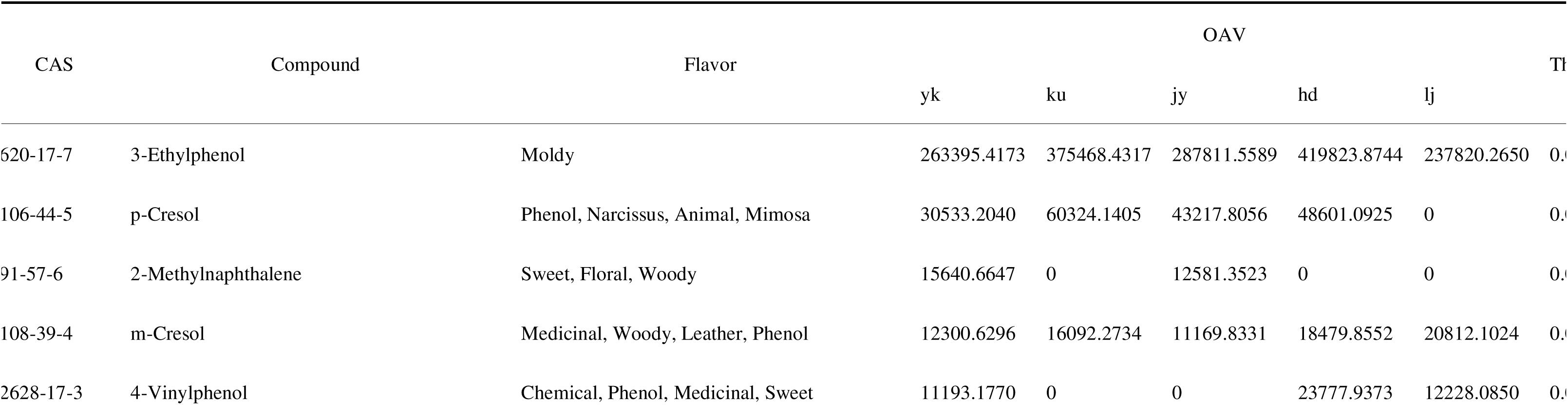
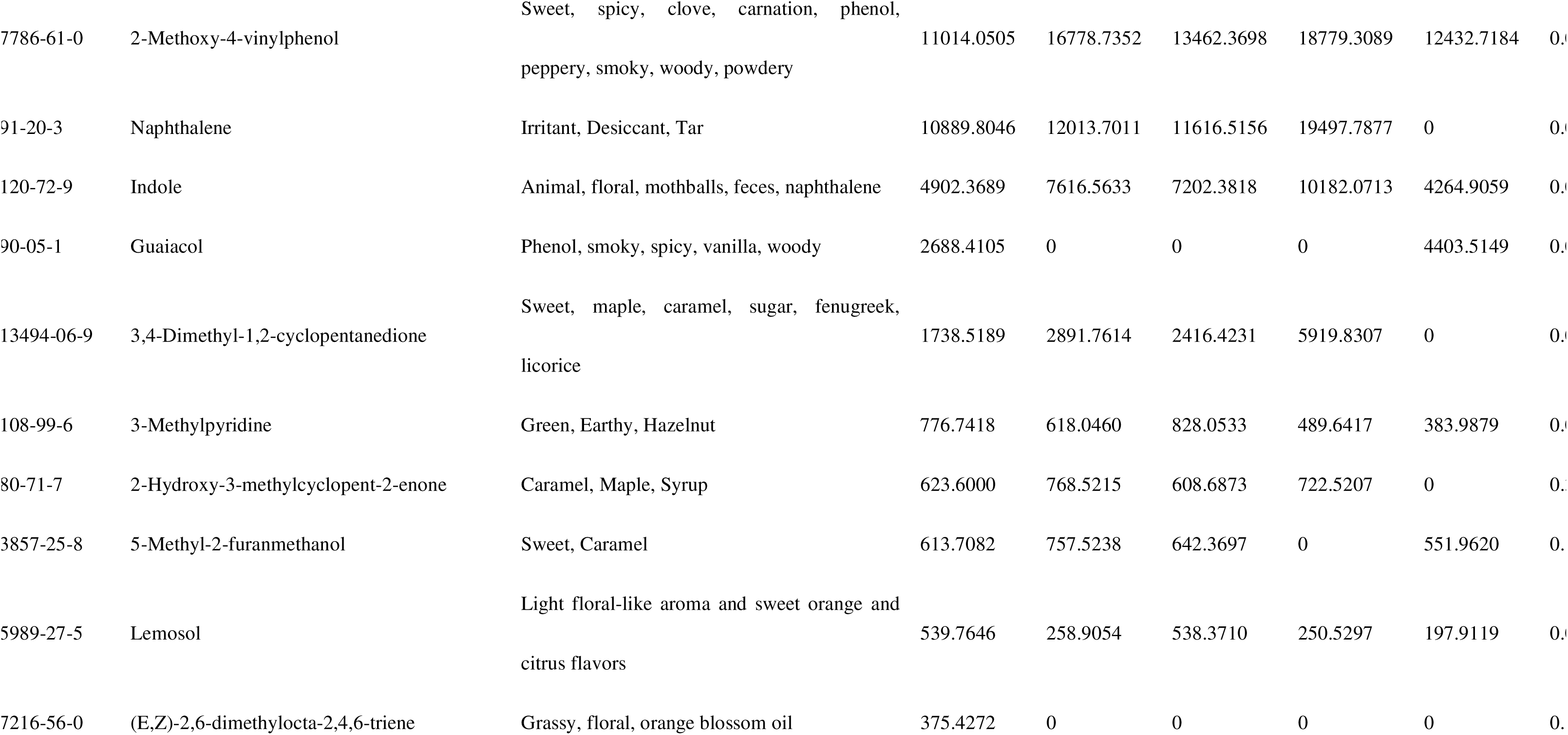
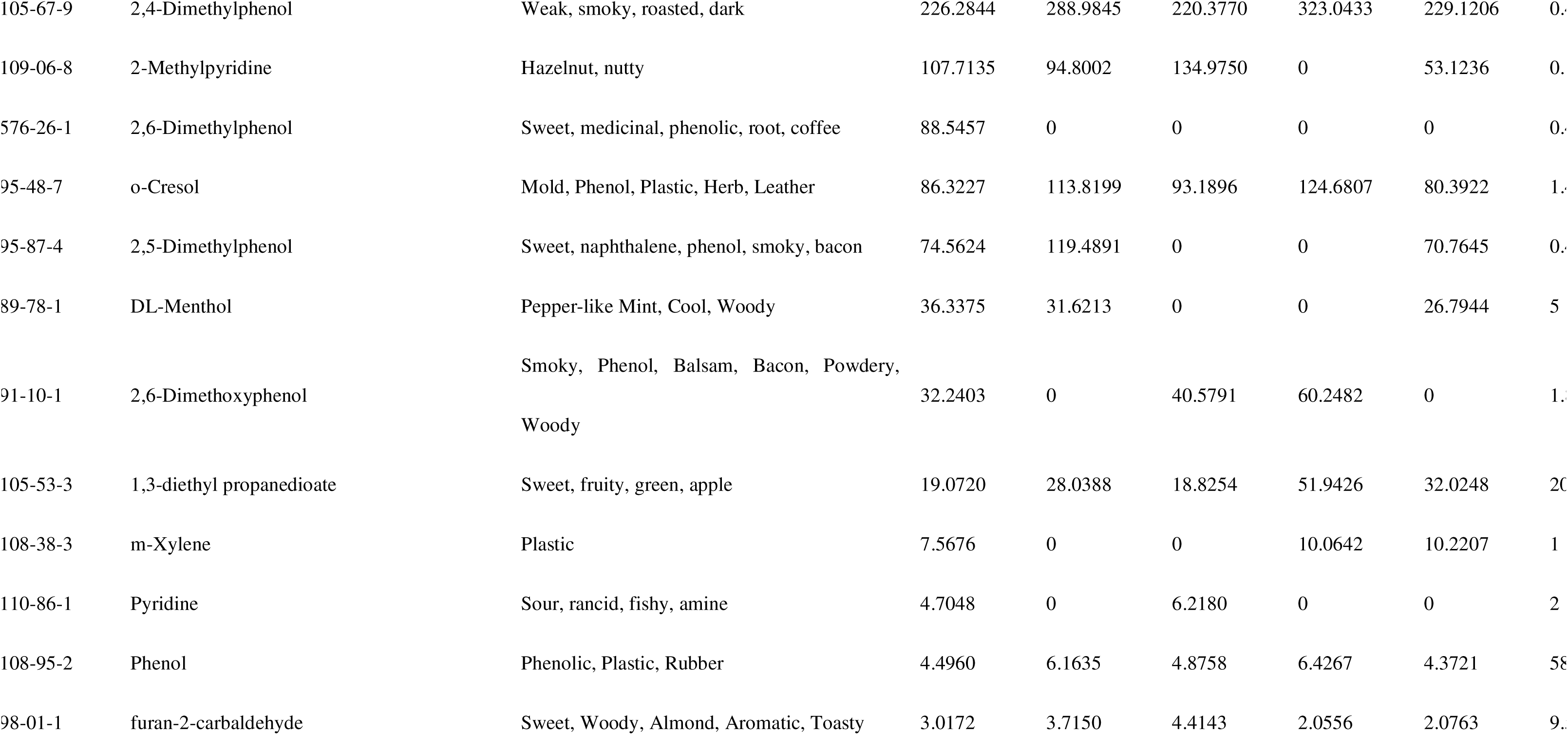
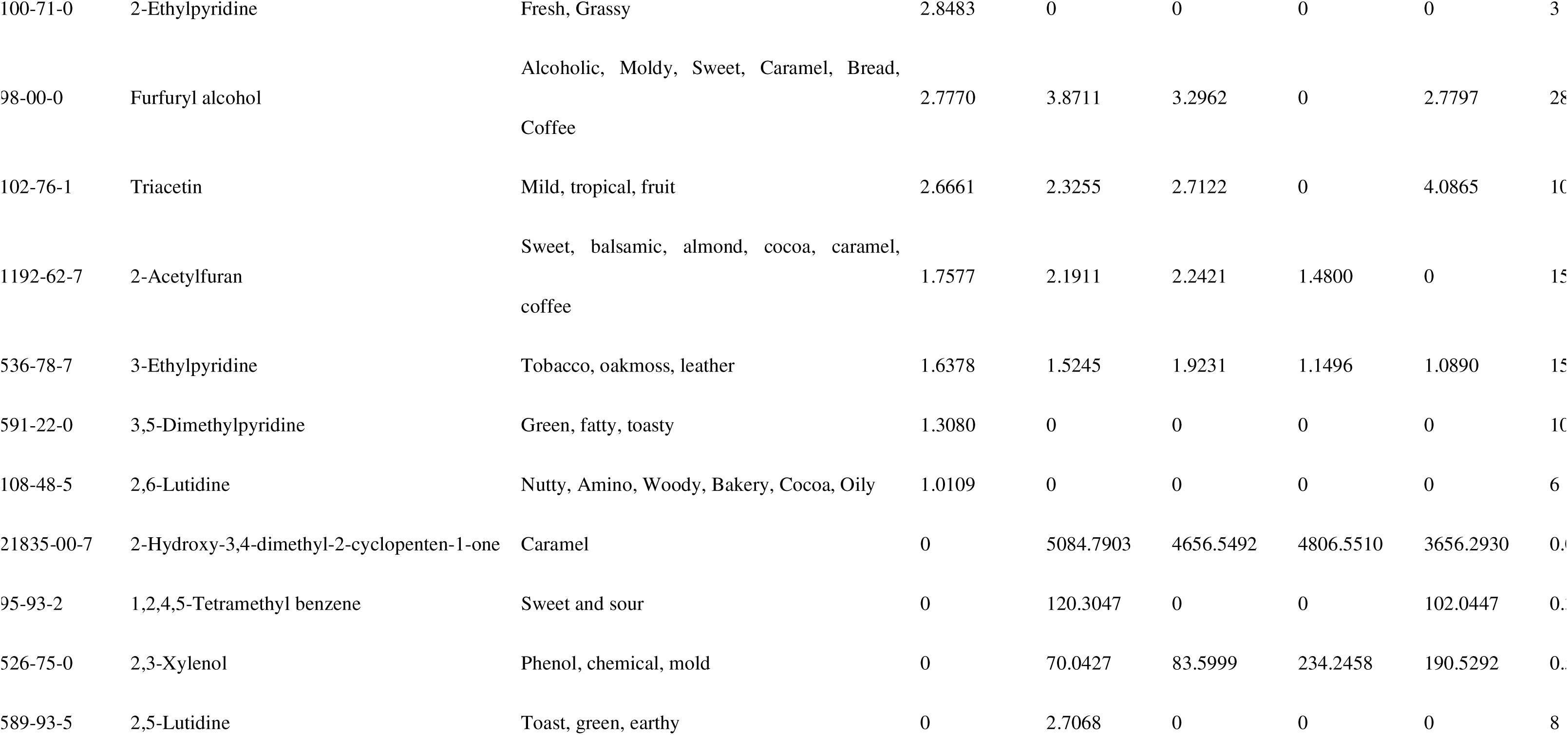
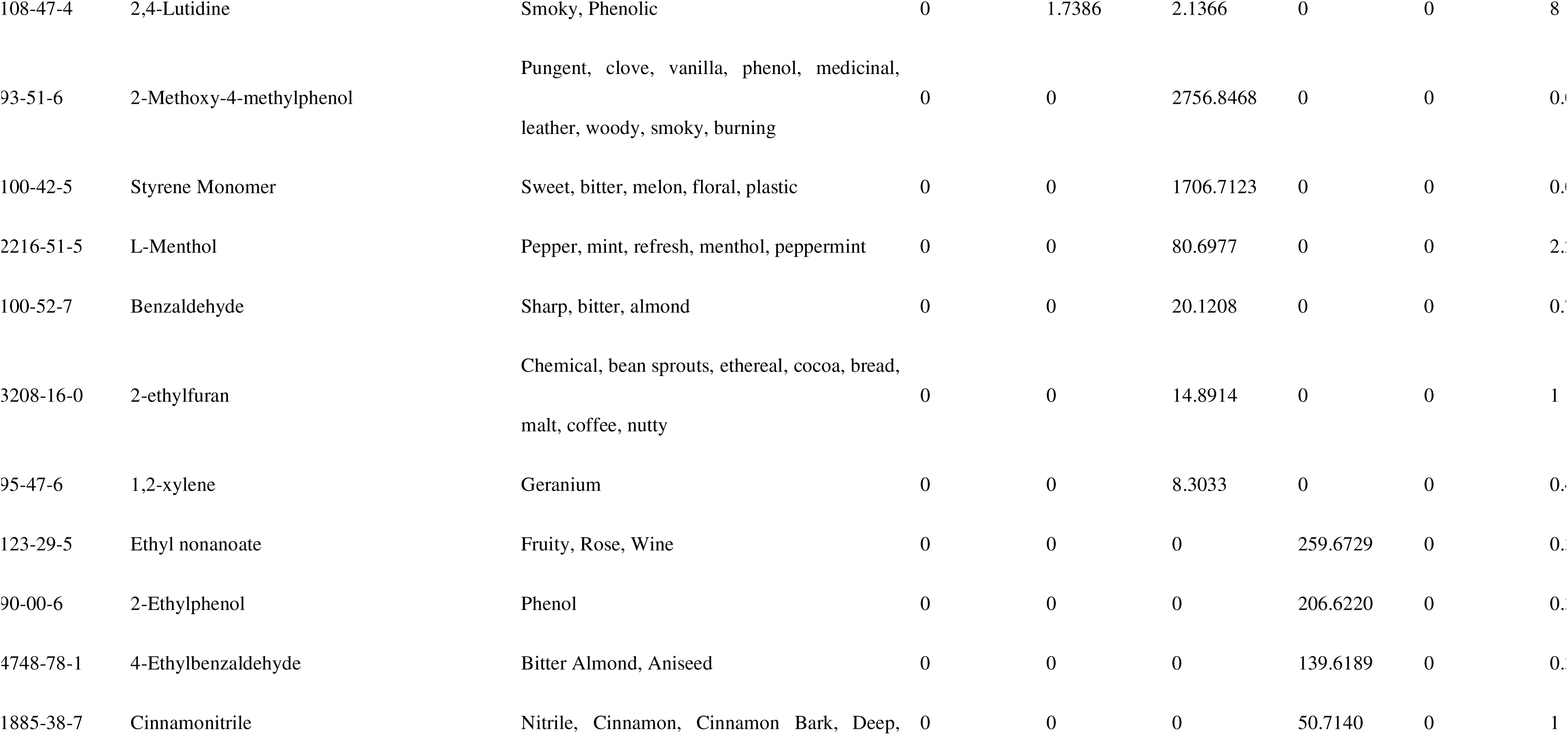
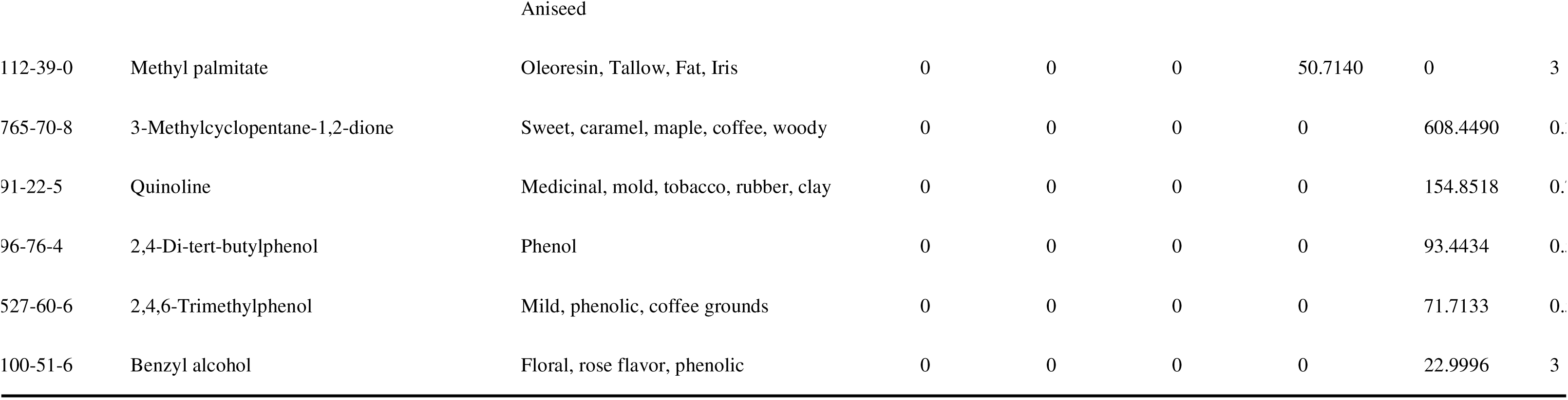
OAV for 5 cigarettes using the safe pre-treatment method.

Among the five different types of cigarettes, we found a total of 55 aroma compounds with OAVs greater than or equal to 1. Of these, 11 compounds, such as 3-Ethylphenol, m-Cresol, 2-Methoxy-4-vinylphenol, and Indole, contribute to the aroma of all five types of cigarettes. There are 22 aroma compounds that are characteristic compounds of individual cigarettes and are present in only one type of cigarette. Furthermore, we observed that eight compounds had OAVs exceeding 10,000 in cigarettes. These compounds include 3-Ethylphenol, p-Cresol, 2-Methylnaphthalene, m-Cresol, 4-Vinylphenol, 2-Methoxy-4-vinylphenol, Naphthalene, and Indole. These substances are considered crucial aroma-contributing compounds, exerting the most significant impact on the differences in aroma among the five types of cigarettes.

Thus, through OAV analysis, we can identify which compounds are most important for contributing to the aroma of different types of cigarettes, providing valuable insights for further research into cigarette aroma characteristics.

From Figure 3, it can be seen that PC1 accounts for 37.8%, and PC2 accounts for 31.7% of the variance, together covering almost 70% of the total information in the data. Based on the results above, it can be inferred that after PCA analysis of the odor activity values of aroma compounds in the five types of cigarettes, the aromas of yk, jy, and ku are relatively similar, whereas they differ significantly from the aromas of hd and lj.

**figure 3.**
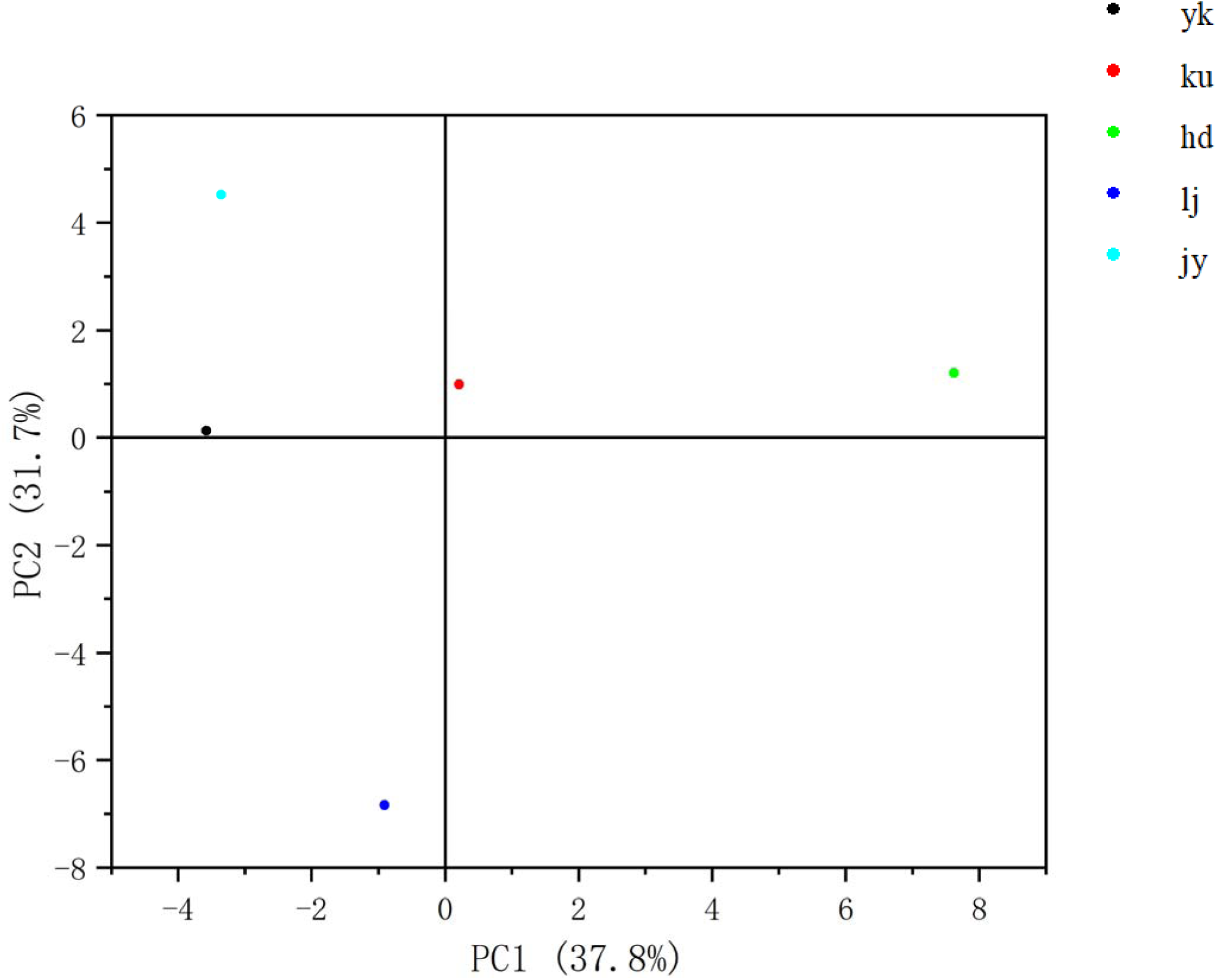
PCA analysis of OAV for five cigarette types using the SAFE pre-treatment method

### Cigarette comfort evaluation results

In this experiment, the smoking comfort evaluation method of cigarettes in Table 1 was used. Fifty volunteers were recruited, 39 men and 11 women, most of whom were between 18 and 25 years old and had smoked for 1 to 5 years. The results showed that the evaluation scores of different volunteers on cigarettes were quite different, so the analysis of the median score of each cigarette was mainly.

**Table 6.** Results of sensory comfort evaluation of Hengta cigarettes.

| No. | Gender | Age | Smoke for years | Score |  |  |  |  |
| --- | --- | --- | --- | --- | --- | --- | --- | --- |
|  |  |  |  | yk | ku | jy | hd | lj |
| 1 | Male | 18~25 | 1~3 years | 76 | 62 | 60 | 86 | 68 |
| 2 | Male | 18~25 | 1~3 years | 66.8 | 64.5 | 65 | 73.2 | 76 |
| 3 | Male | 18~25 | Less than 1 year | 63 | 35 | 43 | 57 | 51 |
| 4 | Male | 18~25 | 1~3 years | 61 | 57 | 75 | 62 | 57 |
| 5 | Male | 18~25 | 3~5 years | 86 | 87 | 91 | 86 | 86.5 |
| 6 | Male | 18~25 | 1~3 years | 61.5 | 62.5 | 58.5 | 55 | 59 |
| 7 | Male | 18~25 | 5~10 years | 76.5 | 75.2 | 69 | 67.1 | 69.5 |
| 8 | Male | 18~25 | 5~10 years | 56 | 63 | 58 | 60 | 58 |
| 9 | Male | 18~25 | 1~3 years | 59 | 88 | 81 | 79 | 84 |
| 10 | Male | 18~25 | 1~3 years | 85 | 75 | 70 | 75 | 71 |
| 11 | Female | 18~25 | 1~3 years | 82 | 47 | 47 | 63 | 70 |
| 12 | Male | 18~25 | 1~3 years | 65 | 62 | 47.5 | 77 | 55 |
| 13 | Male | 18~25 | 3~5 years | 71 | 75 | 66 | 65 | 66 |
| 14 | Male | 18~25 | 1~3 years | 30 | 30 | 25 | 25 | 25 |
| 15 | Male | 26~30 | 5~10 years | 77 | 86 | 76 | 81 | 90 |
| 16 | Male | 18~25 | 5~10 years | 36.5 | 34 | 31 | 26 | 33 |
| 17 | Female | 18~25 | 1~3 years | 55 | 60 | 56 | 60 | 74 |
| 18 | Male | 18~25 | 1~3 years | 90 | 63 | 76 | 87 | 77 |
| 19 | Male | 18~25 | Less than 1 year | 44 | 65 | 70 | 58 | 75 |
| 20 | Male | 18~25 | 3~5 years | 84 | 76 | 78 | 69 | 75 |
| 21 | Female | 18~25 | 1~3 years | 86 | 58 | 48 | 46 | 60 |
| 22 | Male | 18~25 | 1~3 years | 72 | 81 | 85 | 88 | 75 |
| 23 | Female | 18~25 | 1~3 years | 90 | 96 | 82 | 81 | 72 |
| 24 | Male | 18~25 | Less than 1 year | 54 | 56 | 70 | 75 | 86 |
| 25 | Female | 18~25 | 1~3 years | 85 | 65 | 59 | 56 | 55 |
| 26 | Female | 18~25 | 1~3 years | 83 | 74 | 63 | 58 | 59 |
| 27 | Male | 18~25 | 1~3 years | 38 | 41 | 55 | 72 | 56 |
| 28 | Male | 18~25 | 1~3 years | 73 | 79 | 58 | 55 | 50 |
| 29 | Male | 18~25 | 1~3 years | 49 | 75 | 43 | 52 | 48 |
| 30 | Male | 18~25 | Less than 1 year | 83.7 | 72.4 | 69.1 | 70.9 | 67.7 |
| 31 | Female | 18~25 | Less than 1 year | 29 | 24 | 50 | 47 | 30 |
| 32 | Male | 18~25 | 1~3 years | 27 | 39 | 51 | 47 | 26 |
| 33 | Female | 18~25 | 1~3 years | 82 | 81 | 77 | 83 | 74 |
| 34 | Male | 18~25 | Less than 1 year | 81 | 75 | 71 | 79 | 69 |
| 35 | Female | 18~25 | 3~5 years | 44 | 56 | 57 | 51 | 46 |
| 36 | Male | 18~25 | More than 10 years | 49 | 61 | 53 | 57 | 62 |
| 37 | Male | 18~25 | Less than 1 year | 62 | 28 | 77 | 54 | 29 |
| 38 | Male | 18~25 | 3~5 years | 59 | 66 | 47 | 60 | 58 |
| 39 | Female | 18~25 | 3~5 years | 42 | 39 | 26 | 25 | 24 |
| 40 | Female | 26~30 | 3~5 years | 72 | 38 | 26 | 26 | 25 |
| 41 | Male | 18~25 | Less than 1 year | 69 | 82 | 72 | 72 | 76 |
| 42 | Male | 18~25 | 1~3 years | 28 | 31 | 30 | 23 | 28 |
| 43 | Male | 18~25 | More than 10 years | 77 | 69 | 56 | 62 | 77 |
| 44 | Male | 18~25 | 3~5 years | 92 | 90 | 83 | 80 | 83 |
| 45 | Male | 18~25 | 1~3 years | 54 | 69 | 58 | 64 | 56 |
| 46 | Male | 18~25 | 5~10 years | 68 | 58 | 53 | 65 | 70 |
| 47 | Male | 18~25 | 1~3 years | 78 | 70 | 74 | 84 | 58 |
| 48 | Male | 18~25 | 3~5 years | 71 | 88 | 71 | 73 | 73 |
| 49 | Male | 18~25 | 1~3 years | 89.5 | 91 | 89.5 | 83.5 | 84 |
| 50 | Male | 18~25 | 1~3 years | 55.5 | 70.5 | 75.5 | 74 | 67.5 |
| Average score |  |  |  | 65.36 | 63.802 | 61.442 | 63.494 | 61.284 |
| Median score |  |  |  | 68.5 | 65 | 61.5 | 64.5 | 66.75 |

First of all, the score difference of the five cigarettes is small, whether it is the average score or the median score is only less than 7 points. Among them, yk is considered to be the most comfortable cigarette with 68.5 points, followed by lj66.75 points, ku65 points, hd64.5 points, and finally, jy is considered to be the least comfortable cigarette with only 61.5 points, and the score difference between the previous several cigarettes is also large.

The sensory panel consisted of 50 volunteers, including 39 males and 11 females, aged 18–25 years. While this demographic composition allowed us to focus on a specific segment of smokers, it may limit the generalizability of the results, as smoking preferences and sensory perceptions can vary across different demographic groups. To address this limitation, we performed a subgroup analysis to explore potential differences in sensory evaluation scores based on gender and smoking history.

The results of the subgroup analysis revealed no significant differences in scores between male and female participants. Similarly, no significant differences were observed when participants were stratified by smoking history. These findings suggest that, within the studied population, gender and smoking history did not strongly influence the perceived smoking comfort of the five cigarette types. However, we acknowledge that the narrow age range and gender imbalance may limit the broader applicability of our results. Future studies should aim to include a more diverse demographic sample, encompassing a wider age range and a balanced gender distribution, to further validate these findings.

## Conclusion

The innovation of this study lies in the more effective extraction of aroma compounds produced during cigarette smoking using a smoking machine and Cambridge filter. Subsequently, three different methods were employed for the pretreatment of Cambridge filters, and through GC-MS analysis, it was determined that the combination of Cambridge filters with the Solvent Assisted Flavor Evaporation (SAFE) method is the preferable pretreatment method for extracting cigarette aroma compounds. In addition, aroma compounds that contribute significantly to cigarette aroma were identified by calculating odor activity values (oav) and combining thresholds for the aroma of five types of cigarettes, and differences or similarities between different cigarettes were derived by principal component analysis (PCA). Finally, the comfort difference of different cigarettes was obtained through sensory evaluation.

Based on the results of this experiment, future research could involve adjusting the aroma compounds of different cigarettes. Additionally, by integrating sensory experiments, adjustments to the aroma compounds of cigarettes perceived by volunteers to have poor smoking sensations could be made, aiming to improve their aroma.

## Author contributions

Dengke Li: Investigation; Writing—review and editing; Resources. Yuchen Gu: Writing—original draft; Data curation; Formal analysis; Visualization. Chuntao Zhang: Investigation; Methodology; Resources. Ting Fei: Methodology; Resources. Lichao Ma: Investigation; Resources. Ruoxin Wu: Investigation; Methodology. Liqi Tao: Validation; Software. Zhizhang Tian: Resources. Tao Feng: Conceptualization, Resources, Supervision. All authors have read and agreed to the published version of the manuscript.

## Data availability statement

Data supporting the results of this study can be obtained from the corresponding authors upon reasonable request.

## Informed consent

Written informed consent was obtained from all study participants.

## Ethics statements

The sensory evaluation section of this study involved human participants. The sensory evaluation was approved by the Ethics Committee of Shanghai Institute of Technology (SIT-2023-LL15). All panelists signed informed consent forms and were paid after the sensory evaluation.

## References

1. Rennard, Stephen I. “Cigarette Smoke in Research.” American Journal of Respiratory Cell and Molecular Biology 31, no. 5 (2004): 479–80.

2. Chang, Joanne T., Gabriella M. Anic, Brian L. Rostron, Manju Tanwar, and Cindy M. Chang. “Cigarette Smoking Reduction and Health Risks: A Systematic Review and Meta-Analysis.” Nicotine & Tobacco Research 23, no. 4 (2021): 635–42.

3. Das, Salil K. “Harmful Health Effects of Cigarette Smoking.” Molecular and Cellular Biochemistry 253, no. 1 (2003): 159–65.

4. Dechanet, C., T. Anahory, J. C. Mathieu Daude, X. Quantin, L. Reyftmann, S. Hamamah, B. Hedon, and H. Dechaud. “Effects of Cigarette Smoking on Reproduction.” Human Reproduction Update 17, no. 1 (2011): 76–95.

5. Marti-Aguado, David, Ana Clemente-Sanchez, and Ramon Bataller. “Cigarette Smoking and Liver Diseases.” Journal of Hepatology 77, no. 1 (2022): 191–205.

6. Pasupathi, Palanisamy, Govindaswamy Bakthavathsalam, Y. Yagneswara Rao, and Jawahar Farook. “Cigarette Smoking—Effect of Metabolic Health Risk: A Review.” Diabetes & Metabolic Syndrome: Clinical Research & Reviews 3, no. 2 (2009): 120–27.

7. Sherman, Charles B. “Health Effects of Cigarette Smoking.” Clinics in Chest Medicine 12, no. 4 (1991): 643–58.

8. Sherman, Charles B. “The Health Consequences of Cigarette Smoking: Pulmonary Diseases.” Medical Clinics of North America 76, no. 2 (1992): 355–75.

9. Skurnik, Yair, and Yehuda Shoenfeld. “Health Effects of Cigarette Smoking.” Clinics in Dermatology 16, no. 5 (1998): 545–56.

10. Zare, Samane, Mehdi Nemati, and Yuqing Zheng. “A Systematic Review of Consumer Preference for E-Cigarette Attributes: Flavor, Nicotine Strength, and Type.” PLOS ONE 13, no. 3 (2018): e0194145.

11. Omaiye, Esther E., Wentai Luo, Kevin J. McWhirter, James F. Pankow, and Prue Talbot. “Electronic Cigarette Refill Fluids Sold Worldwide: Flavor Chemical Composition, Toxicity, and Hazard Analysis.” Chemical Research in Toxicology 33, no. 12 (2020): 2972–87.

12. Gades, Mari S., Aleksandra Alcheva, Amy L. Riegelman, and Dorothy K. Hatsukami. “The Role of Nicotine and Flavor in the Abuse Potential and Appeal of Electronic Cigarettes for Adult Current and Former Cigarette and Electronic Cigarette Users: A Systematic Review.” Nicotine & Tobacco Research 24, no. 9 (2022): 1332–43.

13. Du, Ping, Rebecca Bascom, Tongyao Fan, Ankita Sinharoy, Jessica Yingst, Pritish Mondal, and Jonathan Foulds. “Changes in Flavor Preference in a Cohort of Long-Term Electronic Cigarette Users.” Annals of the American Thoracic Society 17, no. 5 (2020): 573–81.

14. Soleimani, Farshid, Sina Dobaradaran, Gabriel E. De-la-Torre, Torsten C. Schmidt, and Reza Saeedi. “Content of Toxic Components of Cigarette, Cigarette Smoke Vs Cigarette Butts: A Comprehensive Systematic Review.” Science of The Total Environment 813 (2022): 152667.

15. Purkis, Stephen W., Valerie Troude, Gerald Duputié, and Christian Tessier. “Limitations in the Characterisation of Cigarette Products Using Different Machine Smoking Regimes.” Regulatory Toxicology and Pharmacology 58, no. 3 (2010): 501–15.

16. Duvareille, Charles, Benoit Beaudry, Marie St-Hilaire, Mathieu Boheimier, Cyril Brunel, Philippe Micheau, and Jean-Paul Praud. “Validation of a New Automatic Smoking Machine to Study the Effects of Cigarette Smoke in Newborn Lambs.” Laboratory Animals 44, no. 4 (2010): 290–97.

17. Jalili, Vahid, Abdullah Barkhordari, and Alireza Ghiasvand. “A Comprehensive Look at Solid-Phase Microextraction Technique: A Review of Reviews.” Microchemical Journal 152 (2020): 104319.

18. Engel, Wolfgang, Wolfgang Bahr, and P. Schieberle. “Solvent Assisted Flavour Evaporation - a New and Versatile Technique for the Careful and Direct Isolation of Aroma Compounds from Complex Food Matrices.” European Food Research and Technology 209, no. 3-4 (1999): 237–41.

19. Chaintreau, Alain. “Simultaneous Distillation-Extraction: From Birth to Maturity?Review.” Flavour and Fragrance Journal 16, no. 2 (2001): 136–48.

